# Comprehensive profiling of transcriptional regulation in cartilage reveals pathogenesis of osteoarthritis

**DOI:** 10.1101/2024.06.11.598401

**Authors:** Wen Tian, Shan-Shan Dong, Feng Jiang, Jun-Qi Zhang, Chen Wang, Chang-Yi He, Shou-Ye Hu, Ruo-Han Hao, Hui-Miao Song, Hui-Wu Gao, Ke An, Dong-Li Zhu, Zhi Yang, Yan Guo, Tie-Lin Yang

**Author notes:** Correspondence authors: Tie-Lin Yang, Ph.D., Yan Guo, Ph.D. (T.-L.Y.), (Y.G.). These authors contributed equally to this work (Wen Tian, Shan-Shan Dong, Feng Jiang). Conflict of Interest: The authors declare that they have no competing interests.

## Abstract

Cartilage damage is a leading cause of osteoarthritis (OA) etiology, however, the underlying mechanism governing gene expression regulation in this progress is poorly understood. Here, we described a comprehensive profiling of transcriptional regulation of 235 primary human cartilage samples. We identified 3,352 independent significant expression quantitative trait loci (eQTLs) for 3,109 genes. We explored the candidate casual SNP and its underlying regulatory mechanism using our established functional fine-mapping pipeline by integrating the cartilage-specific ATAC-seq data. We identified 117 causal eQTLs that display allele-specific open chromatin (ASoC) and 547 transcription factor binding-disruption (TBD) eQTLs. We conducted cell type-interaction eQTL (ci-eQTL) analyses based on speculated chondrocyte subtype proportions and revealed the regulation relationship of 120 eQTL-gene pairs showed cell type dependency. Further, by integrating with genome-wide association studies (GWASs) data of OA, we nominated 43 candidate effector genes for OA risk loci. We verified that the T allele of the OA risk variant rs11750646 increased the AR binding affinity to an open chromatin region and promoted the expression of an OA-related gene PIK3R1. Altogether, our findings provide new insights into the unique regulatory landscape of cartilage and elucidate potential mechanisms underlying the OA pathogenesis.

## Introduction

Osteoarthritis (OA) is the most prevalent chronic joint disease, causing pain, disability and loss of function in over 500 million people worldwide(1). OA is mainly caused by progressive degeneration of articular cartilage in joints such as the knee, hip or hand(1, 2). Figuring out the role of articular cartilage in OA and the specific mechanisms of gene expression regulation involved is essential for developing efficient therapies.

Genome-wide association studies (GWASs) have reported more than 100 susceptibility loci for OA(3–9), however, the majority of SNPs map to noncoding regions and the downstream effector genes in articular cartilage have not been well clarified. Expression quantitative trait loci (eQTLs) data provide a valuable tool for screening effector genes of noncoding SNPs. The first eQTL regulation landscape of OA articular cartilage has been constructed from 99 European patients(10), and identified 1,453 genes with eQTLs, which allows the nomination of four effector genes of OA GWAS loci. However, the power to detect mechanistically informative expression signals rely on sufficient numbers of samples from relevant tissues(11), a larger sample size of articular cartilage is needed to obtain more statistically significant eQTLs. Furthermore, given the discrepancy of allele frequencies in different populations, a cartilage eQTL resource from non-European population will provide additional information to elucidate the gene expression roles of GWAS variants. Therefore, it is necessary to conduct eQTL analysis for cartilage in a relatively large non-European population.

Considering the majority of eQTLs are resident in non-coding regions and span hundreds or even thousands of genetic variants, a significant challenge remains in pinpointing the specific variant that is causal within an eQTL. Statistical methods utilizing summary data as input have been developed to refine a whole significant locus to a smaller set for further investigation(12, 13), such as FINEMAP(14), DAP-G(15), SuSiE(16). However, these methods do not decipher the biological effect caused by allele changes. An explicable method is to evaluate the causal variation’s impact on epigenetic features, such as changes in chromatin accessibility and the binding affinity of transcriptional factors (TFs)(17, 18), as important ways to affect gene expression that have been widely reported by us and others(19–22).

Previous studies have shown that some of the eQTL effects are cell type-specific(23–27). Chondrocytes have several subtypes(28–31), but their genetic regulatory heterogeneity remains poorly understood. To address this issue, an approach based on in silico predicted cell subtype proportions from bulk RNA-seq data has been developed to reveal the cell type dependency of eQTL effects(32). This method has been widely applied to bulk RNA-seq data from several tissues(33–36). However, its application specifically to chondrocyte subtypes remains unexplored and represents a critical gap in our understanding of genetic regulatory mechanisms in cartilage.

In this study, we performed comprehensive characterization for transcriptional regulation of knee cartilage samples from 204 OA patients (Fig. 1). We identified *cis*-eQTLs and evaluated the contribution of chondrocyte subtypes to eQTL effects. Integrating our eQTL data with functional fine-mapping results based on our established, along with OA GWAS data, we nominated effector genes of OA risk loci, and revealed the causal variants regulate the effector gene expression through chromatin accessibility alteration or TF binding site disruption. Our study provides a valuable eQTL resource in cartilage and offers new insights into the genetic contributions to OA pathogenesis.

**Figure 1.**
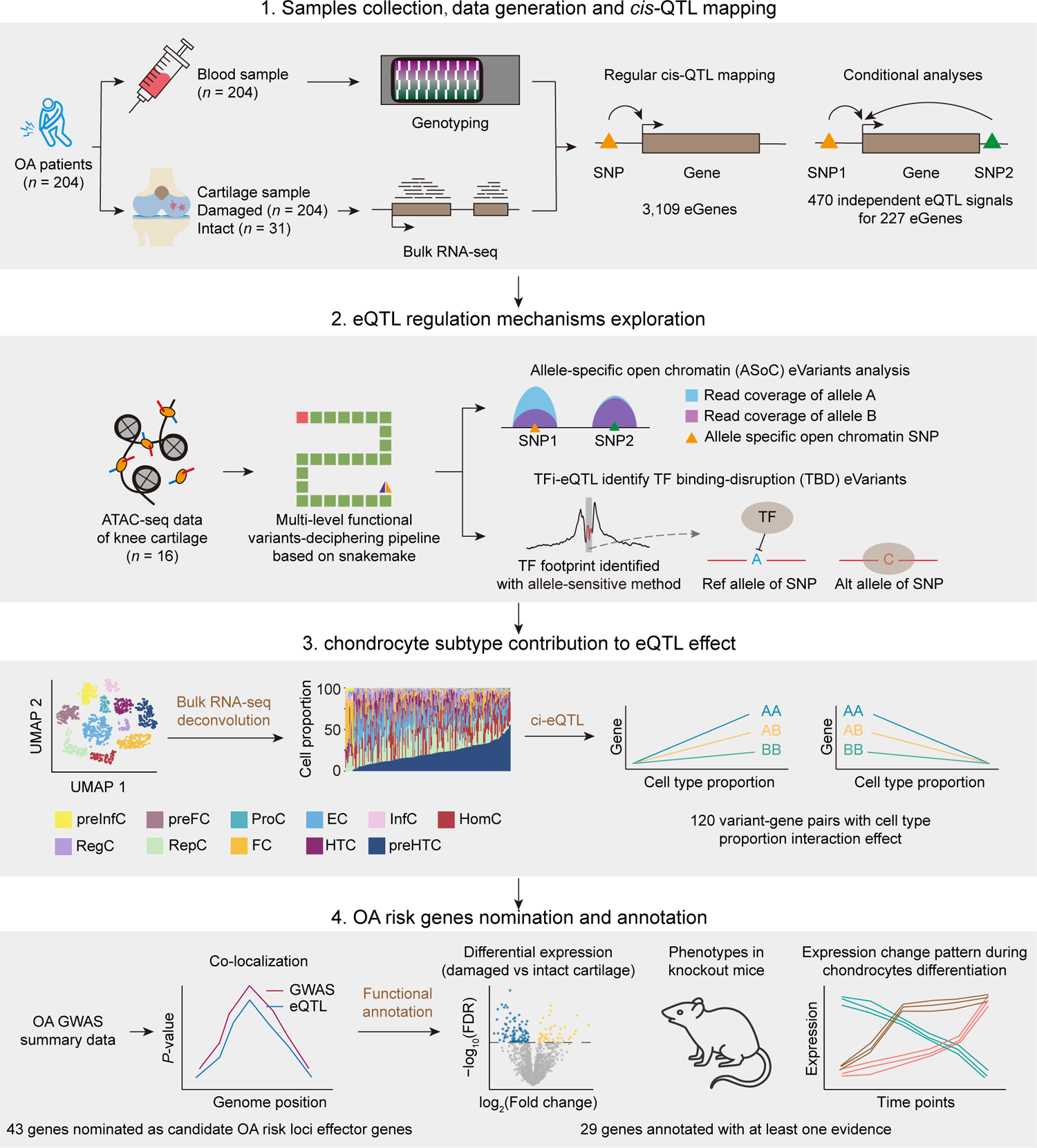
Overview of the study. We collected damaged and intact knee cartilage samples for bulk RNA-seq and blood samples for DNA genotyping. We performed *cis*-eQTL mapping and followed by conditional analysis to identify independent eQTL signals for each eGene. Next, using our established multi-level functional variants-deciphering pipeline, we performed functional fine-mapping based on cartilage epigenetic features that called from cartilage ATAC-seq data to nominate the candidate causal eVariants which may regulate gene expression through alteration of chromatin accessibility and TF binding. We then deconvoluted bulk gene expression data into chondrocyte subtype proportions based on single-cell RNA-seq data of knee cartilage samples and performed cell type-interaction eQTL analyses to reveal the genetically regulatory heterogeneity of gene expression between chondrocyte subtypes. Finally, by integrating our eQTL data and OA GWAS summary data, we nominated OA risk loci effector genes and annotated their function using multiple lines of evidences.

## Results

### *Cis*-eQTL mapping

#### Regular cis-eQTL

We generated 235 qualified knee cartilage RNA sequencing (RNA-seq) data from 204 Chinese Han individuals undergoing joint replacement for OA with extensive quality controls (Fig. 1 and Supplementary Tables 1,2). We obtained a median of 14,809 expressed protein coding genes, 3,407 long non-coding RNA (lncRNA) genes and 1,915 genes of other types (expressed gene was defined as TPM ≥ 0.1, Supplementary Fig. 1a). Most of protein-coding genes were high expressed (TPM ≥ 10), while the majority of non-coding genes had low expressions (TPM < 1) (Supplementary Fig. 1b). Additionally, we generated matched genome-wide genotype data from peripheral blood samples of all donors (Supplementary Tables 1,3). The genotype data showed no significant stratification (Supplementary Fig. 2a), and the minor allele frequencies (MAF) were strongly correlated with that of Asians and weaker correlated with that of Europeans in the 1000 Genomes Project (Supplementary Fig. 2b.c). We then performed *cis*-eQTL (±1 Mb window centered on the transcription start site (TSS)) mapping using RNA-seq data from damaged cartilage. We identified 3,109 genes (eGenes) with at least one genome variant (eVariant) significantly associated with their expression at 5% false discovery rate (FDR) level (Fig. 2a and Supplementary Table 4), including 2,539 (81.7%) protein-coding genes, 404 (13.0%) lncRNAs and 166 (5.3%) of other types (Fig. 2b and Supplementary Table 5). Compared with non-eGenes, eGenes showed significantly higher expression (*P* < 2.2×10^-16^; Wilcoxon rank sum test; Fig. 2c) and lower expression variation coefficient (*P* < 2.2×10^-16^; Wilcoxon rank sum test; Fig. 2c).

**Figure 2.**
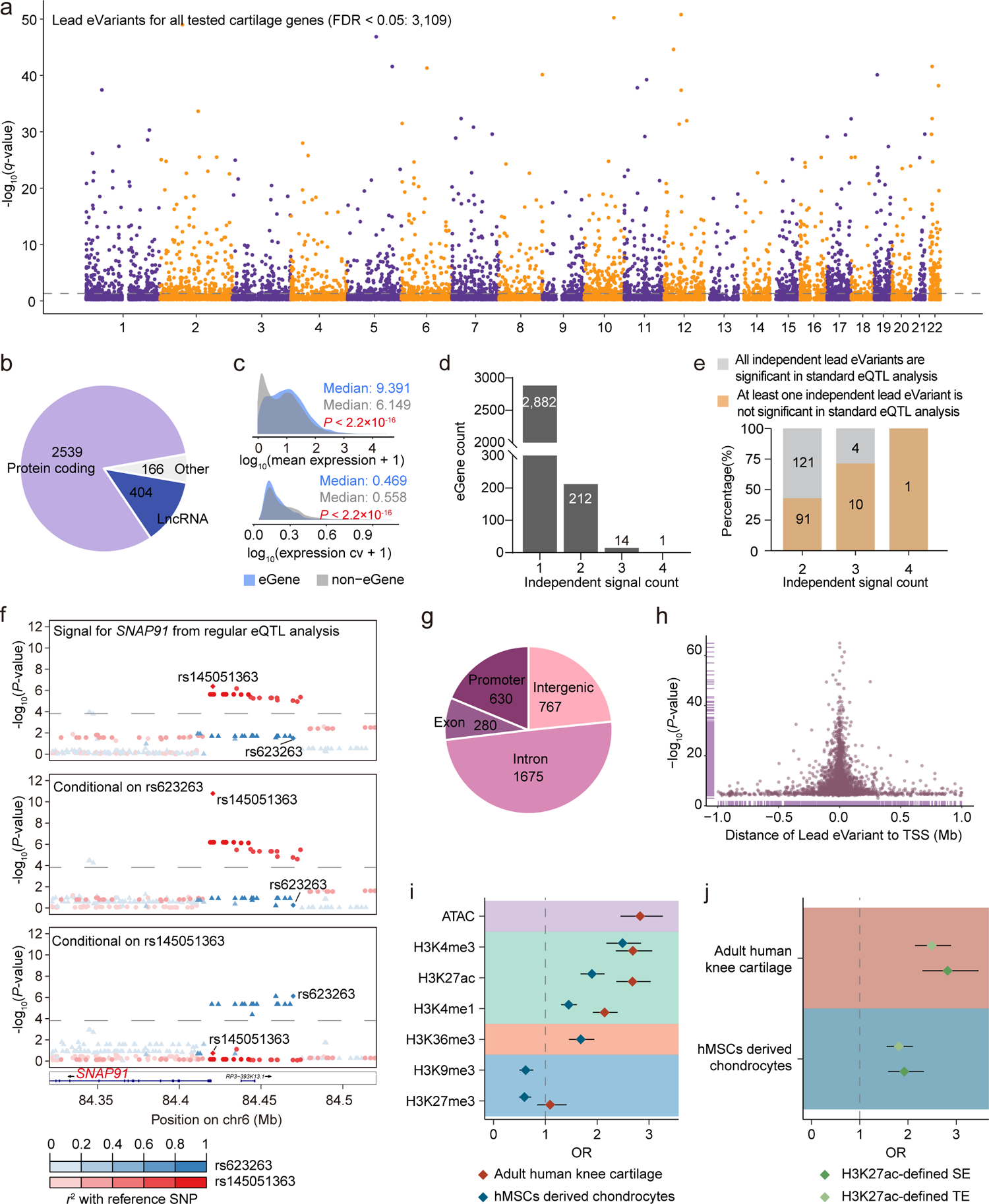
*cis*-eQTL results. **a**, Manhattan plot shows the association strength of lead variants of all eQTL signals of all tested gene (*n* = 18,756). The y axis is the statistical significance in -log_10_(*P*-value). **b**, Biotype distribution of eGenes. **c**, Comparison of expression and variable coefficients distribution of significant eGenes and non-eGenes. The x axis is log-converted mean expression (top panel) and variable coefficients (bottom panel). *P*-value was calculated using Wilcoxon rank sum test. **d** Number of genes with different number of independent signals in conditional analysis. **e**, Distribution of genes with or without newly identified independent signal (not significant in regular *cis*-eQTL analysis) after conditional analysis. **f**, *SNAP91* has two independent eQTL signals tagged by rs145051363 (middle) and rs623263 (bottom), respectively. In regular *cis*-eQTL analysis, only the rs145051363 tagged signal is significant, the rs623263 tagged signal is not significant (top). **g**, Genome distribution of lead eVariants of all independent eQTL signals. **h**, Distance of lead variants of all independent eQTL signals from the transcription start site (TSS) of the associated eGenes and corresponding nominal statistical significance in -log_10_(*P*-value). **i**, Enrichment of LD-pruned (*r*^2^ < 0.8) lead eVariants in different epigenetic features. Data are showed as odds ratio (OR) ± 95% confidence intervals. The utilized epigenetic feature data are from both original adult human knee cartilage samples and hMSCs derived chondrocytes. ATAC-seq data is absent from hMSCs derived chondrocytes. H3K36me3 and H3K9me3 ChIP-seq data are absent from adult human knee cartilage. **j**, Enrichment of LD-pruned (*r*^2^ < 0.8) lead eVariants in H3K27ac-defined SEs and TEs from both adult human knee cartilage and hMSCs derived chondrocytes.

#### Conditional analysis

To identify independent eQTL signals, we conducted conditional analysis and found that 227 of 3.109 eGenes had more than one independent eQTL signals (Fig. 2d and Supplementary Table 6). Importantly, 102 eGenes had at least one newly identified independent signal that could not be detected in regular eQTL analysis (Fig. 2e). For example, rs623263 showed significant association with *SNAP91* expression only when conditional on the effect of rs145051363 which tagged the other independent signal (Fig. 2f). Compared to the lead eVariants (the most significant eVariant for each eGene) already significant in regular cis-eQTL analysis, the newly identified lead eVariants in conditional analysis showed significantly lower MAF (*P* = 0.0282; Wilcoxon rank sum test; Supplementary Fig. 3a), but had no significant difference in distance to the transcription start sites (TSS) of the target genes (*P* = 0.296; Wilcoxon rank sum test; Supplementary Fig. 3b) and effect size for the target genes (*P* = 0.902; Wilcoxon rank sum test; Supplementary Fig. 3c). Downsampling analysis demonstrated our larger sample size contributed to the identification of independent association signals (Supplementary Fig. 4).

For the 3,352 unique independent lead eVariants, the largest part (1,675, 50.0%) was located in the intron regions, followed by the intergenic regions (767, 22.9%), promoter regions (630, 18.8%), and exon regions (280, 8.4%) (Fig. 2g). Lead eVariants were almost symmetrically concentrated on both sides of the TSS of their target genes (Fig. 2h). In total, 2.042 (61.0%) lead eVariants were located within ± 50 Kb from the TSS. In addition, there were 791 lead eVariants (23.6%) located > 100 Kb away from the TSS of the target genes, suggesting long-range regulatory models between eVariants and their target genes.

To assess eQTL reproducibility, we compared our results with a previously published cartilage study involving 99 European individuals(10). Despite the high direction consistency of regulatory effect of the shared variant-gene pairs, our data also uncovered a large number of specific pairs (Supplementary Fig. 5). This was partly attributable to the larger sample size of our study (Supplementary Fig. 6). Additionally, the difference in allele frequency between the Chinese and European populations was a considerable factor. Among the 245,488 specific eVariants identified in our study, 13,432(5.70%) and 4,227 (1.79%) had a MAF < 0.01 or < 0.001 in the 1000 Genomes Project European population, respectively (Supplementary Fig. 7). This meant that if the minimum minor allele count was set to 10, as used in this study, it would require approximately 1,000 and 10,000 samples to calculate their effects on the expression of *cis*-genes, respectively. These sample sizes exceed the maximum sample size of any single tissue in the GTEx project (Muscle-Skeletal, 803 samples, v8)(37), posing a significant challenge in terms of sample collection and highlighting the values of eQTL mapping in different populations.

#### Characteristics of eVariants

We explored the functional characteristics of the eVariants using epigenomic data from human knee cartilage(38) and human mesenchymal stem cells (hMSCs) derived chondrocytes(39). As expected, eVariants were significantly enriched in open chromatin regions marked with ATAC-seq peaks and multiple active histone markers (H3K4me3, H3K27ac, H3K4me1 and H3K36me3) (Fig. 2i). Specifically, eVariants exhibited slightly stronger enrichment in H3K27ac-defined super-enhancers (SEs) than that in typical enhancers (TEs) (Fig. 2j), suggesting a more profound role of SEs in genetic regulation of gene expression.

### Functional fine-mapping reveal candidate causal variants and mechanisms underlying cartilage gene expression

To fill the mechanistic gap between the eVariants and their target genes expression, we established a multi-level functional variants-deciphering pipeline to evaluate whether they could alter the chromatin accessibility of regulatory elements or disrupt the TF binding to the regulatory elements.

#### Allele-specific open chromatin (ASoC) eVariants

We first explored the impact of eVariants on the chromatin accessibility (Fig. 3a). Using the read coverage of heterozygous sites called from knee cartilage ATAC-seq data(20, 40) (Supplementary Fig. 8; Methods), we identified 1,170 allele-specific open chromatin (ASoC) variants showing significant imbalance between the two alleles (nominal *P* ≤ 0.05) (Supplementary Fig. 9 and Supplementary Table 7). A total of 117 eVariants were annotated as ASoC variants and they associated with the expression of 137 eGenes (Fig. 3b and Supplementary Table 7). For example, the C allele of rs3771239 was associated with higher *SDC1* expression and higher local chromatin accessibility around it (Fig. 3c-e). Furthermore, the G allele of rs12471240 was associated with higher *NGEF* expression and higher local chromatin accessibility around it (Supplementary Fig. 10). These results suggesting the potential causal role of rs3771239 and rs12471240 in regulating their target gene expression by affecting the regulatory elements accessibility.

**Figure 3.**
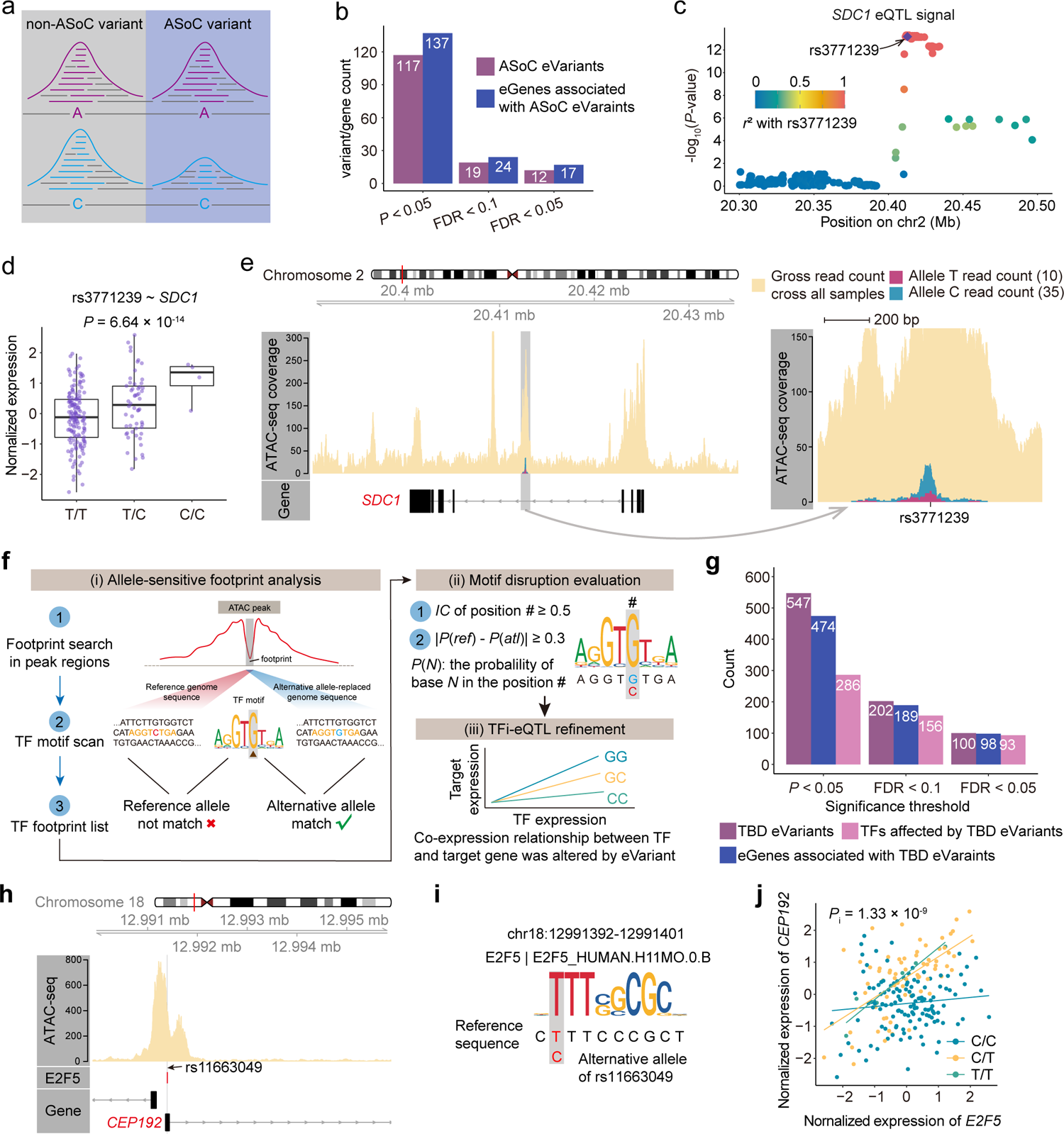
Functional fine-mapping reveal candidate causal variants and mechanisms underlying cartilage gene expression. **a**, The schematic diagram of an allele specific open chromatin variant. **b**, Number of eVariant that is ASoC variant and eGenes associated with ASoC eVaraints under different statistical significance threshold. **c**, Plot show eQTL signal associated to *SDC1*. The y axis is the statistical significance of cartilage eQTL for *SDC1* gene in -log_10_(*P*-value). The x axis is the chromosome coordinate. **d**, Association between the genotypes of variant rs3771239 and *SDC1* gene expression. The y axis is the normalized expression of *SDC1* in damaged cartilage samples; the x axis is the genotype at the rs3771239. The whiskers extend to the 5th and 95th percentiles; Each dot is a different sample. The *P*-value was reported by FastQTL nominal procedure. **e**, Variant rs3771239 was located within an open chromatin region marked by an ATAC-seq peak in the intron region of the *SLC44A2* and altered the local chromatin accessibility. The sequencing reads with C allele piled up a significantly higher peak than that piled up by A allele-contained reads. **f**, Schematic diagram for TF binding-disruption analysis module in our multiple-level functional variants-deciphering pipeline. Three main steps included. For the allele-sensitive TF footprint analysis step, we introduced both the original reference genome sequence and the allele-replaced genome sequence corresponding to the signal valley regions as inputs for TF motif search. In this example scenario, only the alternative allele-replaced genome sequence matched to the TF motif. **g**, The number of eVariant that is TBD variant, eGenes associated with TBD eVaraints and TFs whose binding sites were affected under different statistical significance threshold. **h**, rs11663049 overlapped with a E2F5 footprint. **i**, Alternative allele C at rs11663049 disrupts the motif of E2F5. The motif coordinate was shown on the top. **j**, TFi-eQTL analysis reveal rs11663049 modified the regulatory role of E2F5 on *CEP192* expression. The y axis is the normalized expression of *CEP192* in damaged cartilage samples; The x axis is the normalized expression of *E2F5* in damaged cartilage samples. The *P*-value indicates the statistical significance of interaction term of TF expression and target expression in linear-mixed regression model.

#### TF binding-disruption (TBD) eVariants

We and others had revealed that genetic variants could disrupt the TF binding on regulatory elements, such as enhancer and promoters, and the consequently alter the gene expression(22, 41–45). We further investigated the effect of eVariants on the TF binding. Due to the lack of ChIP-seq data of TFs in cartilage tissues, we established an allele-sensitive TF footprint analysis method, which considered the TF binding specificity to both the reference allele and alternative allele of each variant in motif scan step (Fig. 3f; Methods) to speculate the binding sites for 441 expressed TFs (median TPM ≥ 1) in our damaged cartilage samples (Supplementary Fig. 11a and Supplementary Table 8). We identified 222 to 39,416 non-overlapped unique binding sites for each TF (Supplementary Fig. 11b and Supplementary Table 8). Compared with the traditional footprint analysis, our allele sensitive method could detect a total of 22,627 new footprints for all 673 motifs used for analysis (Supplementary Fig. 12). We validated the accuracy of the footprint results using the CTCF ChIP-seq data from cartilage samples, and found high validation rates for all five motifs of CTCF (Supplementary Fig. 13), indicating the reliability of our method.

Next, we used a TF motif information content-based method to investigate the potential motif disruption effects of eVariants (Fig. 3f; Methods). A total of 1,313 variants were defined as candidate TBD variants in this step (Supplementary Table 9). Further, we performed a TF expression-interaction eQTL analysis (TFi-eQTL) to assess whether the TF for which the motif was disrupted by a variant was the true mediator of the eQTL effect (Fig. 3f; Methods). Briefly, we constructed a linear-mixed regression model with an interaction term of genotype dosage and TF expression to detect the effect of the eVariant on the co-expression of TF and target gene. As a result, we identified 547 TBD eVariants that could regulate the expression of 474 eGenes through the disruption of binding sites of 286 TFs with a nominal significance threshold (Fig. 3g and Supplementary Table 10). Taking the significant regulatory pair rs11663049∼*CEP192* as an example (Supplementary Fig. 14), we found rs11663049 located within a E2F5 footprint (Fig. 3h), and the T allele contributed to the formation of a sequence context matched the E2F5 motif while the C allele disrupted the motif (Fig. 3i). In our TFi-eQTL analysis, samples with T allele showed a stronger positive expression correlation between *E2F5* and *CEP192* (Fig. 3j), suggesting that the expression promoting effect of allele T was mediated by E2F5 binding.

### Evaluation of chondrocyte subtype contribution for eQTL effects

Genetic regulation of gene expression exhibited a broad cellular specificity(23–27). We performed cell type-interaction eQTL analyses (ci-eQTL) to reveal the driver or non-driver cell types for the significant eQTLs in regular eQTL results (Fig. 4a; Methods). We first got deconvoluted proportion for 11 chondrocyte subtypes using signature matrix produced from the latest knee cartilage single-cell RNA-seq data(31) (Fig. 4b and Supplementary Table 11). Using pseudo-bulk expression matrices as validation sets (Methods), we observed high correlation between ground truth and predicted cell proportion deconvolution results using the signature matrix (Supplementary Fig. 15), demonstrating the reliability of deconvolution result. Then the ci-eQTL analysis were applied to the five chondrocyte subtypes (preHTC, RepC, HomC, EC, preFC) which presented largest proportion variance across all samples (Supplementary Fig. 16). We identified 16 to 30 genes with significant ci-eQTLs for different cell types (Supplementary Tables 12-16). Most variant-gene regulatory pairs were aligned to the driver model (Fig. 4c).

**Figure 4.**
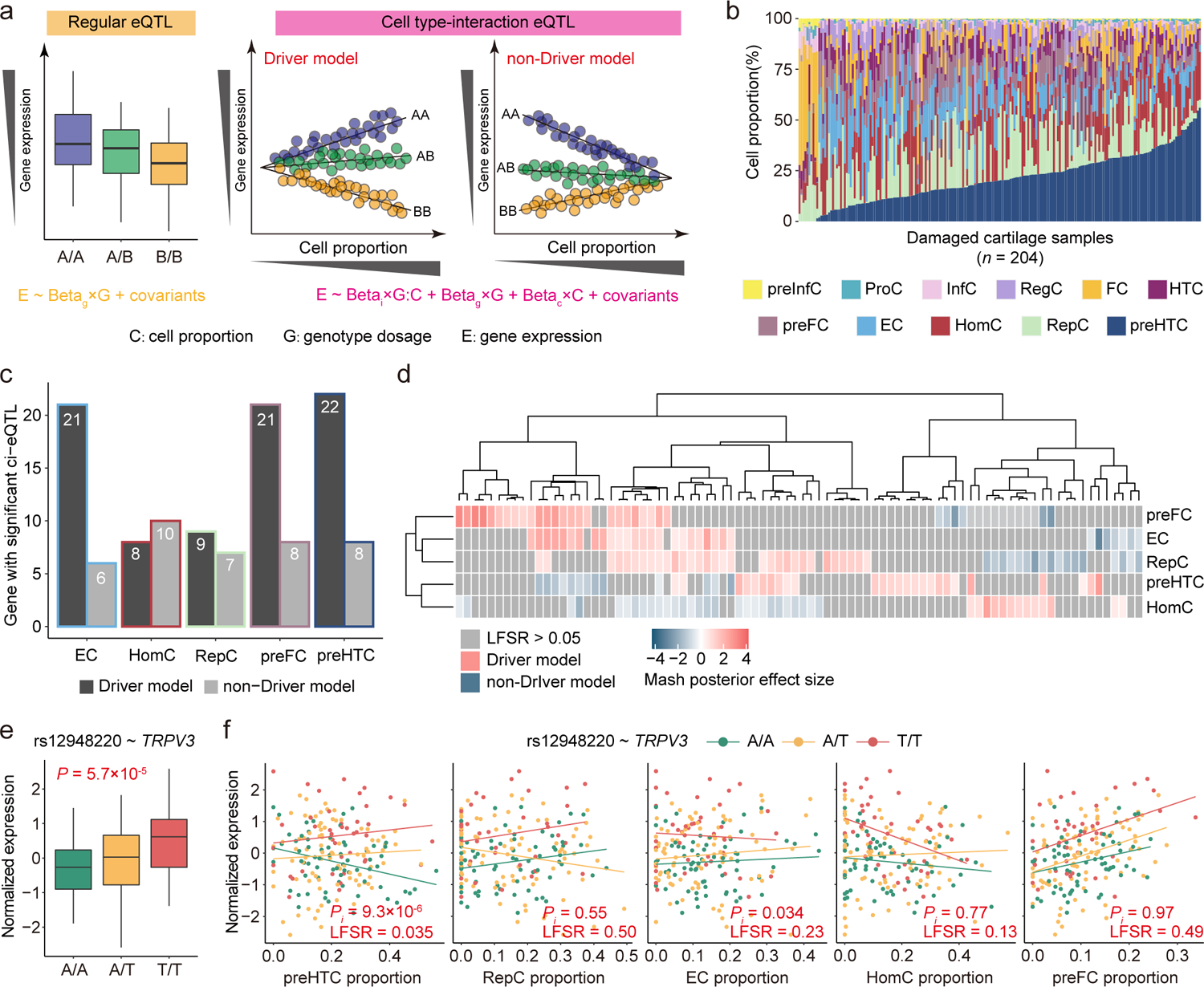
Genetically regulatory heterogeneity between chondrocyte subtypes. **a**, Schematic outline of ci-eQTL analysis. For significant variant-gene pairs in regular *cis*-eQTL analysis, an interaction term of genotype dosage and cell type proportion is modeled in the linear-mixed model to test the cell type dependency of certain eQTL effect. Driver model means the eQTL effect emerging with increase of cell proportion. Non-driver model means that eQTL effect disappearing with increase of cell proportion. **b.** Chondrocytes subtype proportions predicted by CIBERSORTx per sample. Each column is a different sample. The y axis is the accumulated cell proportions. **c**, Number of significant genes discovered in ci-eQTL analyses that aligned to driver and non-driver model for five chondrocyte subtypes. **d**, Heatmap shows the detailed mash results of all significant ci-eQTL results. The color depth represents the mash-reported posterior effect size estimates of interaction item of genotype dosage and cell proportion. Red represents the diver model; Blue represents non-Driver model. Each column is a variant-gene pair. The rows and columns were clustered based on the similarity. **e**, Association between the genotypes of variant rs12948220 and *TRPV3* gene expression. The y axis is the normalized expression of *TRPV3* in damaged cartilage samples; the x axis is the genotype at the rs12948220. The whiskers extend to the 5th and 95th percentiles. The *P*-value was reported by FastQTL nominal procedure. **f**, ci-eQTL results of rs12948220∼*TRPV3* pair across five different chondrocyte subtypes. The y axis is the normalized expression of *TRPV3* in damaged cartilage samples; the x axis is the cell proportion. Each point represents one sample and colored by genotype types. The *P*-values indicate the statistical significance of interaction term in linear-mixed regression model. LFSR was estimated by mash.

We estimated the sharing and specificity of all 120 significant cell type-interaction variant-gene pairs using multivariate adaptive shrinkage (mash) analysis(46). We obtained 86 variant-gene pairs with local false sign rate (LFSR) < 0.05 in at least one cell type (Figure 4d and Supplementary Table 17). Among which, 49 variant-gene pairs assigned as significant driver-model in just one cell type, suggesting potential cell-type specific eQTL effect. For example, the expression of the *TRPV3* showed a significantly positive correlation with the count of T allele at rs12948220 (Fig. 4e). Consistently, with the proportion of preHTC increased, T allele carriers exhibited significantly higher *TRPV3* expression, whereas no such pattern was observed in the other four cell types (Fig. 4f).

### eQTL-GWAS integration nominates risk genes for OA

GWASs have successfully discovered numerous genetic loci associated with OA, however, the underlying gene expression regulatory mechanisms in cartilage remain elusive. Here, we performed co-localization analysis to integrate our cartilage eQTL data (regular, conditional and ci-eQTL results) with 28 publicly available GWAS summary statistics of OA (Supplementary Table 18)(4–9). We identified 43 genes for which the corresponding eQTL signals showed strong evidence (posterior probability of hypothesis 4 (PP4) ≥ 0.7, Methods) that shared a common causal variant with GWAS signals (Fig. 5a, Supplementary Fig. 17 and Supplementary Table 19). Specifically, our eQTL data aided to identified 19 novel effector genes (gene symbols were shown in red in Fig. 5b) of OA risk loci, which could not be nominated by eQTL data of any tissues in the GTEx project and previous cartilage eQTL data (Fig. 5b and Supplementary Figs. 18,19).

**Figure 5.**
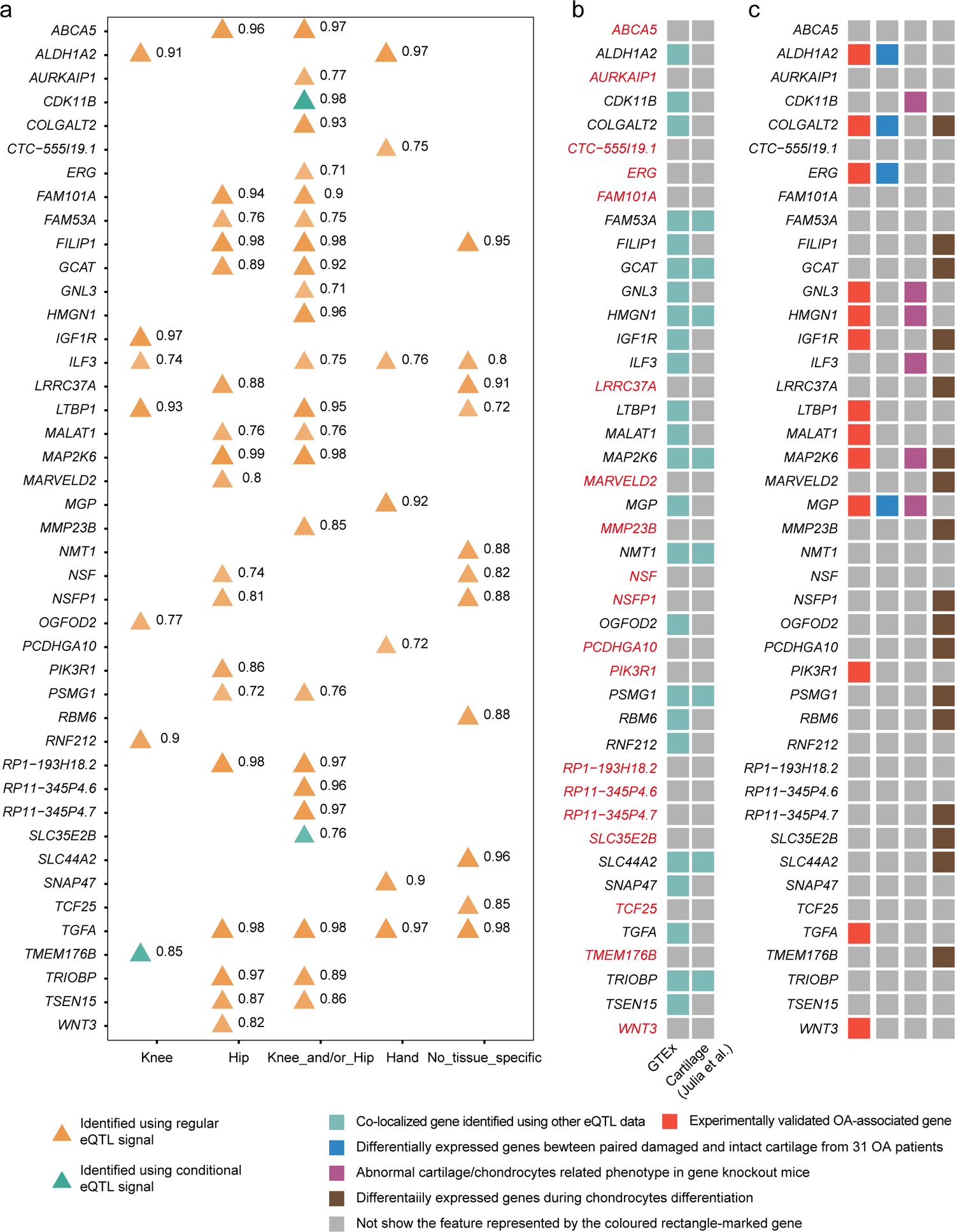
Co-localization analysis nominate risk genes for osteoarthritis. **a**, Co-localization result of OA GWAS loci with our cartilage eQTLs. The numbers next to the triangles represent the highest value of probability of a single shared variant for both GWAS and eQTL signal (PP4) for each type of OA site (see more details in Supplementary Table 13). Genes nominated with PP4 ≥ 0.7 in at least one GWAS data were showed. **b,** Heatmap show whether OA risk loci effector genes nominated using our eQTL data could be nominate by GTEx eQTL data and previous cartilage eQTL data. **c**, Function annotation of co-localized genes from multiple lines of evidences.

Our eQTL data linked OA GWAS signals to 13 OA related genes with experimental validation, four genes showed differential expression between paired damaged and intact cartilage in our data, and six genes showed abnormal cartilage/chondrocytes related phenotypes in gene knockout mice (Mouse Genome Informatics database) (Fig. 5c and Supplementary Tables 20,21). For example, *COLGALT2* showed a significant lower expression in our damaged cartilage samples (log_2_FC = −1.61, *P* = 6.19 × 10^-4^, adjusted *P* = 0.023; Supplementary Table 21). and previous study has reported that *COLGALT2* initiates collagen glycosylation to organize and stabilizes collagen triple helix(47). Particularly, our eQTL data linked OA GWAS signals to three experimentally validated OA genes (*ERG*, *PIK3R1* and *WNT3*) that could not be linked by eQTL data of GTEx tissues and previously reported cartilage tissue(10). For example, *ERG* is a member of ETS transcription factor, the overexpression of ERG inhibited the expression of *MMP1*3 (ref.(48)), which breaks down type II collagen leads to the degeneration of extracellular matrix in the process of OA(49).

To further elucidate the potential functions of our nominated OA risk loci effector genes in the process of chondrocytes differentiation, we generated gene sets from a time series expression data of chondrocytes differentiation(50). Briefly, we classified all differentially expressed genes into 12 clusters to show the overall expression tendency during the chondrocyte differentiation (Supplementary Fig. 20a and Supplementary Table 22). Among these clusters, we identified 17 co-localized genes belonged to these gene clusters (Fig. 5c and Supplementary Fig. 20). We performed gene ontology (GO) enrichment analyses to reveal the function of each cluster, and identified four clusters (cluster 3, 6, 7 and 12) enriched in biological processes related to chondrocytes differentiation (Supplementary Fig. 20b). We found five nominated OA risk effector genes (*FILIP1*, *OGFOD2*, *MMP23B*, *SLC44A2* and *COLGALT2*) belonged to these four clusters (Fig 5c and Supplementary Fig. 20a), suggesting their potential role in the cartilage differentiation. Additionally, for six genes (*NSFP1*, *TMEM176B*, *PCDHGA10, MAP2K6*, *MARVELD2* and *SLC35E2B*) from luster 9 and 10, despite the top enriched biological processes of the gene clusters they belonged to were not directly associated with cartilage formation, their expressions were gradually decreased during chondrocyte differentiation (Supplementary Fig. 20a), suggesting their negative effect on chondrocytes differentiation. Taken together, leveraging our tissue match eQTL data, we provided a high confidence OA risk loci effector gene list for future investigations.

### *SLC44A2* expression might be regulated by OA risk variant rs3810154 via altering chromatin accessibility

The *SLC44A2* was nominated as an OA risk locus effector gene by co-localization analyses with our eQTL data (Fig. 6a) and it showed upregulated expression during chondrocytes differentiation, suggesting a candidate role in the cartilage formation (Fig. 6b and Supplementary Fig. 20). The rs3810154 was located within the co-localized region and the G allele promoted the expression of *SLC44A2* (Fig. 6a, c). Intriguingly, we found the rs3810154 was located in an open region downstream *SLC44A2* and the local chromatin region around the G allele at rs3810154 showed significantly higher accessibility than that of A allele (44 reads for G allele; 20 reads for A; *P* = 3.69 × 10^-3^; binomial test; Fig. 6d), suggesting that rs3810154 regulated *SLC44A2* expression by altering the chromatin accessibility.

**Figure 6.**
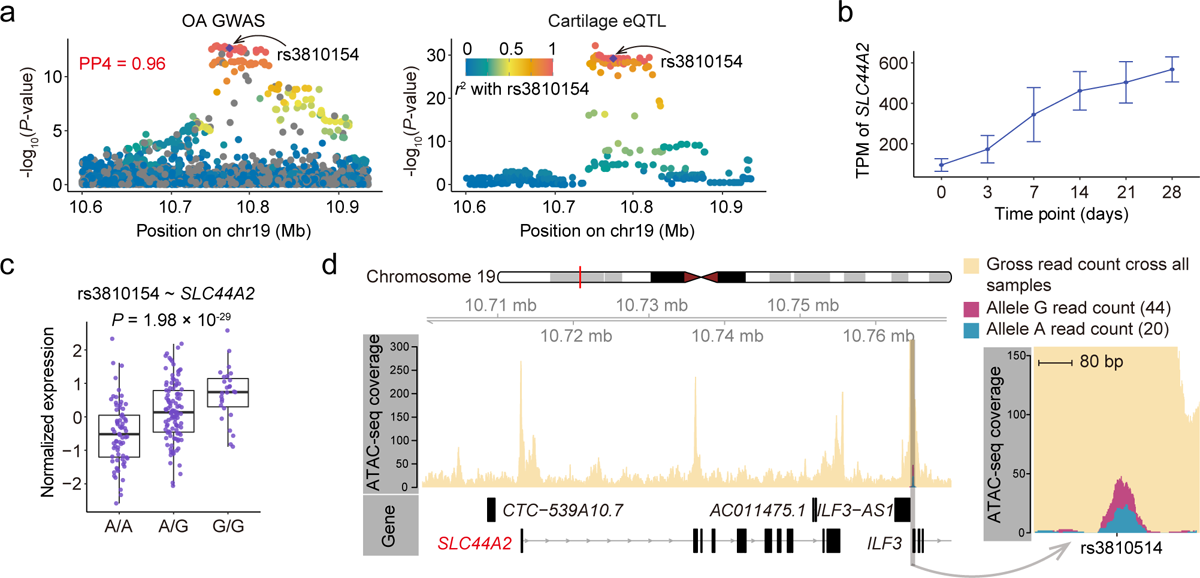
*SLC44A2* expression might be regulated by OA risk variant rs3810154 via altering chromatin accessibility. **a**, Co-localization of OA GWAS locus and eQTL signal associated to *SLC44A2*. The y axis is the statistical significance of OA GWAS and cartilage eQTL for *SLC44A2* gene in -log_10_(*P*-value). The x axis is the chromosome coordinate. **b**, *SLC44A2* expression at each time point during the chondrocyte differentiation. The y axis is the expression value in TPM; The x axis is the time point. Each time point has three biological replications; The data are showed as mean ± sd. **c**, Association between the genotypes of variant rs3810154 and *SLC44A2* gene expression. The y axis is the normalized expression of *SLC44A2* in damaged cartilage samples; the x axis is the genotype at the rs3810154. The whiskers extend to the 5th and 95th percentiles; Each dot is a different sample. The *P*-value was reported by FastQTL nominal procedure. **d**, Variant rs3810154 was located within an open chromatin region marked by an ATAC-seq peak downstream the *SLC44A2* and altered the local chromatin accessibility. The sequencing reads with G allele piled up a significantly higher peak than that piled up by A allele-contained reads.

### *PIK3R1* expression was regulated by OA risk variant rs11750646 via affecting AR binding

*PIK3R1* was nominated as an OA risk locus effector gene only through our eQTL data (Fig 7a and Fig. 5b). It encodes the regulatory subunit 1 of phosphoinositide-3-kinase (PI3K), which inhibits the chondrocyte apoptosis via PI3K/Akt pathway and thus might act as a protected factor against the OA development(51). The rs11750646 was located within the co-localized region (Fig. 7a). The T allele significantly increased the expression of *PIK3R1* (*P* = 2.54 × 10^-8^; Fig. 7b) and disrupted the AR motif in an open chromatin region downstream the *PIK3R1* (Fig. 7c,d). TFi-eQTL results showed that rs11750646 significantly altered the co-expression relationship between *AR* and *PIK3R1* (*P* = 4.43 × 10^-3^_;_ Fig. 7e), indicating that *AR* was the regulator affected by rs11750646.

**Figure 7.**
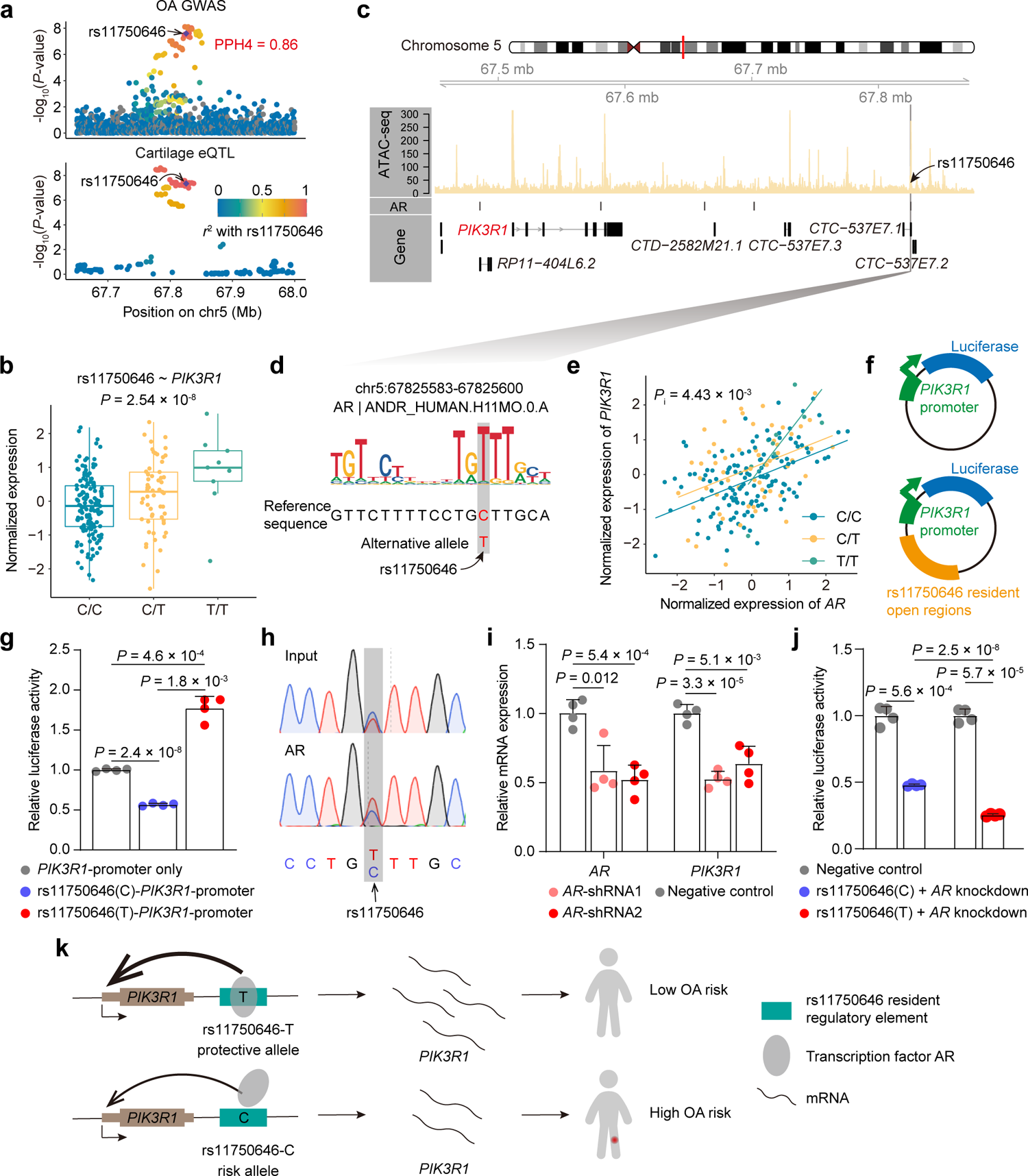
*PIK3R1* expression was regulated by OA risk variant rs11750646 via affecting AR binding. **a**, Co-localization of OA GWAS locus and eQTL signal associated to *PIK3R1*. The y axis is the statistical significance of OA GWAS and cartilage eQTL for *PIK3R1* gene in -log_10_(*P*-value). The x axis is the chromosome coordinate. **b**, Association between the genotypes of variant rs11750646 and *PIK3R1* gene expression. The y axis is the normalized expression of *PIK3R1* in damaged cartilage samples; the x axis is the genotype at the rs11750646. The whiskers extend to the 5th and 95th percentiles; Each dot is a different sample. The *P*-value was reported by FastQTL nominal procedure. **c**, rs11750646 overlapped with an AR footprint that was identified in an open chromatin region marked by an ATAC peak downstream the *PIK3R1*. **d**, Reference allele C at rs11750646 disrupts the motif (ID: ANDR_HUMAN.H11MO.0.A) of AR. The motif coordinate was shown on the top. The symbol (+) represents the genome sequence shown is positive strand. **e**, TFi-eQTL analysis reveal rs11750646 modified the regulatory role of *AR* on *PIK3R1* expression. The y axis is the normalized expression of *PIK3R1* in damaged cartilage samples; The x axis is the normalized expression of *AR* in damaged cartilage samples. The *P*-value indicates the statistical significance of interaction term of TF expression and target expression in linear-mixed regression model. **f**, Schematic diagram of dual luciferase reporter vector. **g**, Detection of activity effect on open chromatin region containing rs11750646 (T and C allele) compared with pGL3-*PIK3R1*-promoter in SW1353 cells by dual-luciferase reporter assay. Luciferase signal was computed as the ratio of firefly luciferase activity to Renilla signal and relative activity was normalized by pGL3-*PIK3R1*-promoter. *P*-values are calculated using two-tailed Student’s *t*-test. **h**, AR preferred the binding to the T allele at rs11750646 confirmed by ChIP followed by Sanger sequencing in SW1353 cells. **i**, Effect of *AR* knockdown on *PIK3R1* expression in SW1353 cells. Two independent shRNAs (shRNA1 and shRNA2) and shRNA-NC (negative control) were used. RT-qPCR analysis of gene expression normalized to *GAPDH* expression level. *P*-values are calculated using two-tailed Student’s *t*-test. **j,** Dual-luciferase reporter assay containing rs11750646-C or -T allele plasmids co-transfected with *AR* shRNAs in SW1353 cells. The shRNA-NC is used as the negative control. Luciferase signals are normalized to Renilla signals. *P*-values are calculated using two-tailed Student’s *t*-test. **k**, The OA risk allele C at rs11750646 disrupts binding of AR to a regulatory element, resulting in decreased *PIK3R1* expression and further increasing the risk of osteoarthritis.

We further performed functional experiments to validate our hypothesis. Using human chondrocyte-like cell line SW1353, the dual-luciferase reporter assay revealed that rs11750646-T contained sequence showed significant enhancement ability on target gene expression, while rs11750646-C contained sequence showed slightly inhibiting ability for target gene expression (Fig. 7f,g). Further, in line with in silico motif analysis results, our ChIP-PCR followed by Sanger sequencing results proved that AR preferentially bound to the T allele at rs11750646 *in vivo* (Fig. 7h). Knockdown of AR significantly repressed *PIK3R1* expression (Fig. 7i). Dual-luciferase reporter assay showed that the gene expression-promoting role of rs11750646-T contained fragment was significantly abolished after AR knockdown, while rs11750646-C contained fragment had a relatively weaker change (Fig. 7j). Collectively, our integrated computational and functional results suggested that AR could allele-preferentially bind the protective T allele (Beta = −0.0521 in GWAS) at rs11750646 to reinforce *PI3KR1* expression and thus associated with OA (Fig. 7k). It was noteworthy that the binding-preferred allele for AR at this SNP was T, which represented an alternative allele not presented in human reference genome sequence version hg19 (Fig. 7h). This AR footprint only identified with our allele-sensitive footprint analysis method, highlighting its great value in uncovering biological mechanisms of genomic variants.

## Discussion

Here, we reported the eQTLs in cartilage based on our samples with the largest sample size to date. We did functional fine-mapping and revealed the candidate causal variants and their underlying regulatory mechanism for hundreds eQTLs. We also evaluated the contribution of chondrocytes subtype to the eQTL effets. Finally, by integrating with OA GWAS data, we extended the list of OA risk loci effector genes.

As an efficient approach, eQTL analysis has been widely used to investigate the genetic impact on the gene expression of human tissues or cell types. As for OA research, Julia et al. published the only eQTL so far for articular cartilage, which is the main tissue related to OA(10). In our present study, we doubled the sample size and reported a larger number of significant eQTLs for cartilage samples, and significantly complemented the previous research. We also successfully distinguished the independent eQTL signal from each other for 237 genes and further reported the capacity of conditional analysis to uncover the eQTL signals that have been missed in the regular eQTL analysis, indicating a new value for conditional analysis.

We made efforts to fill the mechanism gap between genomic variants and gene expression with a functional fine-mapping strategy. We revealed that some eVariants were involved in the alteration of chromatin accessibility. Increasing number of studies have reported the chromatin accessibility quantitative trait loci (caQTLs)(21, 52–54), through which could link the GWAS loci to chromatin accessibility regulation and further to gene expression alteration. However, there are no such resources for cartilage samples. As a compromise, we generated an allele-specific open chromatin SNP set using the publicly available ATAC-seq data from cartilage samples. Although it limited the detectable number of variants that affect chromatin accessibility, we still identified more than one hundred ASoC eVariants with nominal *P*-value threshold. We anticipate that future caQTL data in cartilage will enhance the discovery of variants that regulate gene expression by modulating chromatin accessibility.

We also established an allele-sensitive TF footprint analysis method and identified footprints for 441 expressed TFs in cartilage samples. This approach contributed to the discovery of more TF footprints, especially the AR binding site we have verified (Fig. 7), demonstrating the value of our method. We evaluated the disruption effect of eVariants on the TF motif and then used a TFi-eQTL analysis strategy to refine the variant to a subset that significantly affect the co-expression relationship between TFs and their target genes. With a nominal threshold, we identified more than five hundred potential causal eVariants and the corresponding TFs they impacted.

Integration of eQTL and GWAS is a powerful method for unraveling the molecular mechanism that influence complex traits. Tissue non-matched eQTL data from the GTEx project have broadly been used for co-localization analyses to reveal the gene expression regulation role of OA GWAS loci(5, 6). However, the tissue specific genetic regulation effect may lead to inevitable false nomination of true effector genes. While efforts have been made to nominate effector genes using cartilage expression data derived annotation information, limitations still remained. For example, Hollander et al. reported the transcriptome-wide allelic imbalance data in OA cartilage to link the GWAS variants to gene expression(55), however, this approach could only assess variation in exons. Additionally, existing OA studies have limited the nomination of risk genes using co-localization analysis due to relatively small sample size. Here, by integrating our cartilage eQTL data and OA GWAS data, we nominated 19 novel OA GWAS risk loci effector genes that could not be detectable by other studies. Our data presented in this study helped to make a big step in OA pathogenic mechanism analysis. Based on our functional fine-mapping results, we presented two examples to show how the OA risk variants rs3810154 and rs11750646 affect *SLC44A2* and *PIK3R1* expression through chromatin accessibility and AR binding, respectively.

In summary, we performed comprehensive profiling for transcriptional regulation in knee cartilage from OA patients, reported the largest cartilage eQTL dataset to date, and nominated new effector genes for OA GWAS risk loci. We revealed potential causal variants for both OA risk and gene regulation based on epigenetic feature resources identified from cartilage specific data. Our study provides a valuable resource to elucidate regulatory mechanisms for genes underlying OA etiology.

## Materials and Methods

### Sample collection

All patients clinically diagnosed with OA (KL grade ≥ 3) and required total knee replacement surgery from 2019–2022 in Xi’an Honghui Hospital were eligible for this study after signing the informed consent. During total knee replacement surgery, we isolated the knee damaged or/and intact cartilage with scalpels after the knee joint was removed from the donors and froze it in liquid nitrogen rapidly followed by transferred to a −80L refrigerator for storage. Besides, we collected 2-3 ml peripheral blood from the participants. The summary of clinical information for donors, including age, sex was shown in Supplementary Table S1.

### DNA isolation, genotyping and imputation

DNA was extracted from whole blood using FlexGen Blood DNA Kit (CWBIO) according to the manufacturer’s instructions. DNAs were genotyped using Illumina Infinium Asian Screening Array-24 v1.0 (ASA) containing 659,184 variants in total. TOP strand-based genotypes were reported in plink format using GenomeStudio software (Illumina) and then converted to reference strand-based genotypes using PLINK (v1.9). Subsequently, we performed initial quality control at variants and sample level following the pipelines used by the Genotype-Tissue Expression (GTEx) consortium(37) and that introduced in ref,(56) with a little modification. Briefly, at variants level, keeping variants with consistent genome position and allele with that of 1000 Genomes Project, with genotype missing rate across all samples < 0.02, MAF > 0.01 and the Hardy–Weinberg Equilibrium *P* > 1 × 10^-6^. At sample level, keeping samples with genotype missing rate across all variants < 0.02, with matched sex tag between clinical record and genotype-imputed result, with heterozygosity within 3 standard deviations from the mean heterozygosity.

Genotype imputation was performed using IMPUTE2 (v2.3.2)(57) with all 1000 Genome Project genotype data (phase 3) as reference panel. To increase the imputation accuracy, we changed the default values of the parameter -k and -buffer to 100 and 300, respectively. After imputation, we performed another round of quality control, keeping variants with quality score > 0.3, genotype missing rate across all samples < 0.05, MAF > 0.05 and the Hardy-Weinberg Equilibrium *P* > 1 × 10^-6^, keeping samples with genotype missing rate across all variants < 0.05. We performed principal component analysis (PCA) using smartpca(58) to detect the stratification of population.

### Cartilage RNA isolation, sequencing and data processing

Cartilage RNA was extracted with TRIzol. RNA integrity (RIN) was assessed using the Fragment Analyzer 5400 (Agilent Technologies). Samples with RIN ≥ 5.6 were used for sequencing library preparation using NEBNext^®^ Ultra™ RNA Library Prep Kit for Illumina^®^ (NEB) according to manufacturer’s instructions. RNA sequencing was performed on an Illumina Novaseq 6000 platform and 150 bp paired-end reads were generated, approximately 20 million reads produced for each sample.

Clean RNA-seq reads were mapped to human reference genome hg19 (only contained autosomes, sex chromosomes and mitochondrial chromosomes) with STAR aligner (v2.7.9a)(59) based on the GENCODE v19 (ref.(60)) annotation, the parameters setting was same as the GTEx v8 pipeline. Gene-level read counts and TPM values were generated using RNA-SeQC (v2.3.5)(61) with default parameters. The read mapping rate and base mismatch rate were calculated from the output of SAMtools (v1.9)(62) stats sub-command. The proportions of genomic origin of sequencing reads were produced using QualiMap (v2.2.2-dev)(63). Samples are needed to meet the following metrics: read mapping rate ≥ 0.9, base mismatch rate ≤ 0.01, intergenic mapping rate ≤ 0.3, and rRNA mapping rate ≤ 0.3. In this study, all samples passed this mapping quality control step. Then, we conducted expression outlier filtration with the method described in ref.(64). Briefly, we calculated the pairwise expression correlation coefficients using the log transformed read count of all genes, let the *r_ij_* denote the expression correlation coefficient between sample *i* and sample *j*, we calculated

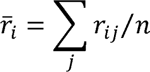

the average correlation coefficient of sample *i* with all others of the total *n* samples. Lower (r_--_ represented a lower quality. Then we calculated

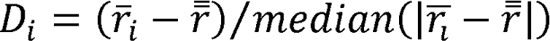

to provide a sense of distance from the grand correlation mean r. Samples with *D* < −5 were considered as outliers and removed. Finally, samples that passed both genotype and RNA-seq quality control were used for the following analysis, which contained a total of 245 samples, of which, 204 were damaged cartilage samples and 31 were intact cartilage samples.

### *Cis*-eQTL mapping

#### Expression data preprocessing

We followed the *cis*-eQTL mapping pipeline from the GTEx consortium(37). Expression data from damaged cartilage samples were used for eQTL analysis. Expression data were normalized using TMM to correct the library size difference between samples, then the rank-based inverse normal transformation (RINT) was applied for each gene across all sample. Finally, genes with TPM > 0.1 and read count ≥ 6 in at least 20% samples were selected. Further, genes on sex and mitochondrial chromosomes were removed.

#### Covariates file generation

PEER factors(65) were calculated to estimate the hidden confounders of gene expression. Following the advice from the GTEx consortium, 30 PEER factors were used for covariates as our sample size is between 150 and 250. Integrated with sex, age and the first two principal components of genotype, we produced a covariate file used for *cis*-eQTL mapping.

#### Regular cis-eQTL mapping

All variants within 1 Mb of transcription start site (TSS) of each gene were used for regular *cis*-eQTL mapping using FastQTL (v2.0)(66) and nominal *P*-value was calculated for each variant-gene pair. Beta distribution-adjusted empirical *P*-values (from adoptive permutation results by setting --permute 1000 10000 for FastQTL) of top variant of each gene were used for multiple testing correction (FDR estimation) using the method described by Storey and Tibshirani(67), which was implemented in R package qvalue, and FDR ≤ 0.05 was used as a threshold to identify genes with significant eQTL (eGenes). To assess the association significance of variants with eGenes, first, we got the mean of Beta distribution-adjusted empirical *P*-values of two genes closest to the 0.05 FDR threshold (one > 0.05 and the other one ≤ 0.05) as genome-wide empirical *P*-value threshold (*p*_t_), then the nominal *P*-value threshold for each gene was calculated using the two beta distribution parameters of the minimum *P*-value distribution obtained from the adoptive permutation procedure for the gene based on the genome-wide empirical *P*-value (F^-1^ (p_t_), F^-1^ denote the inverse cumulative distribution of beta distribution). For each gene, variants with a nominal *P*-value below the gene-level nominal *P*-value threshold were considered statistically significant (eVariants).

#### Conditional analysis

To get the independent *cis*-eQTL signal for each eGene, a two-step conditional analysis (a forward stepwise regression followed by a backward selection) was performed using cis_independent mode of tensorQTL(68) with the regular *cis*-eQTL analysis result as input, the lead variant for each independent eQTL signal was reported. For visualization of each post-conditional independent eQTL signal of single gene, we integrated the genotype dosage of the others lead variants of independent signal into the covariates file and re-conducted the *cis*-eQTL analysis with FastQTL for the gene to get a new *P*-value for each variant.

### Epigenetic data processing

#### ATAC-seq data

The ATAC-seq data of knee cartilage samples from 16 donors were downloaded from GEO database (GSE108301)(40). The reads were mapped to human reference genome hg19 with Bowtie2 (v2.3.5)(69) for each sample, reads with mapping quality < 30 or duplicated were filtered. Peak calling was performed with MACS2 (v2.2.7.1)(70, 71) as described before(42). Peaks presented only in one sample were removed and the others were merged to produce a consistent peak set.

#### Histone modification ChIP-seq data

ChIP-seq data for four histone modifications (H3K4me1, H3K4me3, H3K27ac, and H3K27me3) of adult knee cartilage samples were downloaded from from GEO database (GSE111850)(38). The reads were mapped to human reference genome hg19 with Bowtie2 (v2.3.5) for each sample, reads with mapping quality < 30 or duplicated were removed. Then the bam files from three biological replications of each modification type were merged to a single bam file. Peak calling was performed with MACS2 (v2.2.7.1), for H3K4me1 and H3K27me3, the broad peaks were called with --broad-cutoff set to 0.05, for H3K4me3 and H3K27ac, the narrow peak were called with -q set to 0.05.

Peak files, which were generated by ENCODE project(39) using pseudo-replications-based method, for six histone modification (H3K27ac, H3K27me3, H3K36me3, H3K4me1, H3K4me3, and H3K9me3) of hMSCs derived chondrocytes were download from GEO database under the access number GSE188124, GSE187652, GSE187044, GSE187055, GSE187641, and GSE187471, respectively. Then the genome coordinates were transformed from hg38 to hg19 using liftOver with hg38ToHg19.over.chain.gz annotation file. The bam files of H3K27ac ChIP-seq data were downloaded from ENCODE portal. The alignment coordinates were transformed to hg19 using CrossMap(72) with hg38ToHg19.over.chain.gz annotation file and then the bam files for replication of each modification type were merged.

#### Super-enhancers identification

Super-enhancers identification were based on the H3K27ac ChIP-seq data using ROSE2(73) with default settings. The H3K27ac peaks were used for indicating the enhancer position, and the merged bam files were used for enhancer signal strength estimation.

### eVariants enrichment analyses

All lead variants of conditional independent eQTL signals were LD-pruned with a threshold *r*^2^ < 0.8 to produce a query set. Then for each query variant, we extracted 50 random variants from the SNPsnap dataset(74) as a background set, which accounted for MAF deviation (± 0.05), distance to gene deviation (± 0.5), gene density deviation (± 0.5) and LD buddy deviation (± 0.5). The epigenetic enrichment analysis was performed using fisher’s exact test.

### Identification of allele-specific open chromatin (ASoC) SNPs

We adopted the pipeline described in ref.(20) to identify the variants that showed imbalanced chromatin accessibility with a little modification(Supplementary Fig. 8). Briefly, BAM files from 16 ATAC-seq data of cartilage samples(40) were calibrated by WASP(75) and used as inputs for GATK (v4.2.6.1)(76). Following the GATK Best Practice recommendation, HaplotypeCaller sub-command was used for variants calling based on human hg19 genome sequence and corresponding dbSNP version 138 annotation. Then, the construction and application of InDels and SNPs recalibration model were performed in tandem using VariantRecalibrator and ApplyVQSR sub-command, respectively. In addition, the databases used for VariantRecalibrator as true training set were hg19 versions of Mills and 1000G gold standard InDels list (priority = 12), HapMap v3.3 and 1000G_omni v2.5 (priority = 12), the database used as non-true training set was 1000G phase 1 high confidence SNP list (priority = 10) and dbSNP 138 (priority = 2), the database used as known set was dbSNP 138 (priority = 2). Heterozygous variants with tranche level > 99.0% and located within ATAC-seq peaks were retained. Due to the low read coverage in individual samples, we added up the read depth corresponding to each allele across all heterozygous samples after checking the consistency of allelic effect direction. Finally, we did the binomial test for variant sites with minimum read depth count (DP) ≥ 20 using R function binom.test with a null hypothesis that the two alleles showed the same effect on chromatin accessibility (namely the parameter p was set to 0.5). FDR control was applied using R function p.adjust.

### TF binding-disruption analysis

#### Allele-sensitive TF footprint analysis

Footprint analysis was limited to the expressed TFs which were defined as the genes with at least one DNA binding motif included in JASPAR 2022 core non-redundant set(77) or HOCOMOCO v11 (ref.(78)) database and with a median TPM ≥ 1 across all damaged cartilage samples.

TF footprint analysis involved a step to search TF motifs in the signal valleys in the ATAC-seq peak regions using sequence-match-based method. We supposed there was a possible scenario where the reference genome-stored allele of a SNP may strongly disrupt a TF motif and the alternative allele may help to form a sequence context suitable for TF binding. To account for this, we produced an alternative allele--replaced genome sequence based on hg19 using SNPsplit (v0.6.0)(79) by replacing the reference allele to the alternative allele of all variants presented in genotype data of our samples. Both the original reference genome sequence and newly generated allele-replaced genome sequence were used for TF footprint analysis.

We performed TF footprint analysis using TOBIAS (v0.13.3)(80) with default settings, TOBIAS was specially developed to account for the ATAC-seq data specific bias. All 16 bam files from ATAC-seq pipeline described above were merged to produce a single input to increase the detection power. The search regions were limited to the peaks presented in at least two samples. All motifs corresponding to the expressed TFs were used for motif search. The results reported by TOBIAS were the genome coordinates of the motifs and the speculated binding status.

#### TF binding-disruption evaluation

To investigate the potential impacts of eVariants on the TF binding, we applied a TF motif information content-based method. Briefly, for each motif of expressed TF, we calculated the information content (IC) of each position based on the position frequency matrix (PFM). For each position of the motif, the total possible IC was fixed

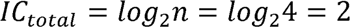

where *n* denote the number of all possible base could be presented in this position, thus *n* is 4 for any TF motif. Then, let the *P(N)* denote the probability of a certain base in the position, we calculated the sum of the uncertainty of all bases (A, C, G, T):

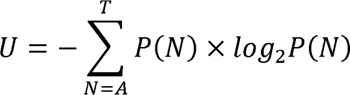

Finally, the final information of the position was calculated:

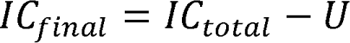

The variants located in a position that with *IC_final_* ≥ 0.5 for a certain motif, and the | P(ref) - *Palt*≥0.3 were considered as candidate TF binding-disruption variants.

#### TF expression-interaction (TFi-eQTL) eQTL analysis

To verify whether the TF for which the motif was disrupted by a variant was the true mediator of the eQTL effect, we performed TFi-eQTL analysis which was similar to the ci-eQTL analysis. Briefly, we modeled following two linear-mixed regression models with lme4 (v1.1-30)(81):

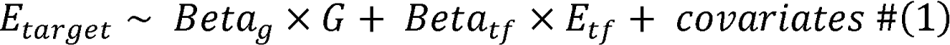

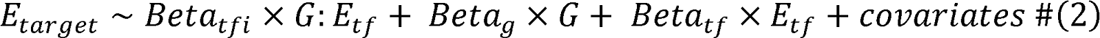

where *E_target_* denote the RINT normalized expression value of target gene. *E_tf_* denote the RINT normalized expression of TF. *Beta* denotes the effect size of each item. Covariates including sex, age, the first two principle components of genotype and 30 peer factors. Sex was modeled as random effect and the other covariates were modeled as fixed effect. The difference of those two models was compared with analysis of variance (ANOVA) implemented by R package pbkrtest (v0.5.1)(82) and a *X*^2^ *P*-value was calculated to indicate the statistical significance. For each TF-target pair for test, the motif and variant with smallest *P*-value was retained.

### Deconvolution of bulk RNA-seq data into chondrocyte subtype composition

Processed single-cell RNA-seq data of knee cartilage(31) was downloaded from the GEO database (GSE255460), which contain 135,896 cells from 19 samples. We randomly selected 5,000 cells from the count matrix, where the columns were named by cell type and the rows were named by gene symbols, as input for CIBERSORTx(83) web server to produce a chondrocyte subtype signature matrix. Then the signature matrix was applied to the TPM matrix of the bulk RNA-seq expression data to predict the proportions of chondrocyte subtypes using the CIBERSORTx web server. S-mode was turned on for CIBERSORTx to correct the technical bias between single-cell and bulk RNA-seq data.

To test the reliability of deconvolution result, we used the following strategy produced a pseudo-bulk expression matrixs as test set: Randomly selecting 5,000 cells from cell list that not used to produce signature matrix each time and repeat for 235 times. this procedure produced a pseudo-bulk matrix contain 235 samples. The CPM for each gene in a pseudo-bulk sample were calculated as the mean value of CPM for this gene across all cells contained in the pseudo-bulk sample. Then we got the predicted cell proportions for the pseudo-bulk samples using the signature matrix and compared the predicted cell proportions with the ground truth.

### Cell type-interaction eQTL (ci-eQTL) analysis

To measure the contribution of a single chondrocyte subtype to eQTL effect, the significant variant-gene pairs in regular *cis*-eQTL result were used for ci-eQTL analysis(35). We modeled following two linear-mixed regression models with R package lme4 (v1.1-30)(81):

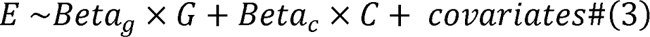

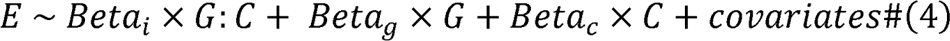

where *E* denotes the RINT normalized gene expression value. *G* denotes the genotype dosage. *C* denotes the proportion of specific chondrocyte subtype (ranging from 0 to 1). *Beta* denotes the effect size of each item. Covariates including sex, age, the first two principle components of genotype, 30 peer factors and proportions of the other 10 chondrocyte subtypes. Sex was modeled as random effect and the other covariates were modeled as fixed effect.

The difference of those two models was compared with analysis of variance (ANOVA) implemented by R package pbkrtest (v0.5.1)(82) and a *X*^2^ *P*-value was calculated to indicate the statistical significance. We extracted the lead variant with the smallest *X*^2^ *P*-value for each tested gene and corrected the *X*^2^ *P*-value with Benjamini-Hochberg FDR control method(84). Genes for which the lead variants with FDR < 0.05 and *P*-value < 5 × 10^-5^ of genotype dosage and cell proportion interaction item were considered as statistically significant cell type-interaction eGenes. We divided the significant variant-gene pairs into driver and non-driver model based on the effect direction of *Beta_g_ Beta_i_* from formula (4). The same effect direction meant that the eQTL effect increased as the cell fraction increased (driver model), suggesting the eQTL effect may come from the cell type used for ci-eQTL analysis (Fig. 4a). In contrast, the opposite effect direction meant that the eQTL effect decreased as the cell fraction increased (non-driver model), suggesting the eQTL effect didn’t come from the cell type used for ci-eQTL analysis (Fig. 4a).

### Sharing and specificity estimation of the ci-eQTL effect

To assess whether the cell type-interaction variant-gene pairs were shared or specific across multiple chondrocytes subtypes (EC, HomC, RepC, preFC and preHTC). We applied the multivariate adaptive shrinkage method (mash)(46), which was implemented in R package mashr (v0.2.57), based on the workflow used for the GTEx data (https://github.com/stephenslab/gtexresults). The effect size of interaction item of genotype dosage and cell proportion (*Beta_i_*) and the standard error (s.e.*_Betai_* calculated form formula 4) were used as inputs for mashr. We extracted the variant with the smallest *P*-value across all three cell types, and all tested variants for each gene and produced two matrices which saved their effect sizes and standard errors for the gene in three cell types, respectively. In addition, two matrices with the same format for a 200,000 randomly selected variant-gene pairs set also be produced. A mash model was built using the exchange effects mode to estimate the prior. The model was then applied to all significant variant-gene pairs for each cell type to get the posterior mash effect sizes and the local false sign rate (LFSR). Pairs with LFSR < 0.5 were considered statistically significant.

### Co-localization analyses

We downloaded a total of 28 OA GWAS summary data from six published studies(4–9) (Supplementary Table 18).For each GWAS summary data, we clumped the genome-wide significant (*P* < 5 × 10^-8^) variants with PLINK parameter “--clump-r2 0.1 --clump-kb 1000” to get the index SNP. The nominal *P*-value for all variants located within ±1 Mb window of the index variant were used for co-localization analysis. For gene tested in cis-eQTL analysis, the nominal *P*-value for all *cis*-variant (± 1 Mb) of each gene were extracted for co-localization analysis. For genes with more than one independent signal, the *P*-values calculated from the conditional analysis were used.

Co-localization analysis was performed using R package coloc (v5.1.0.1)(85), which implemented a Bayesian test to assess whether two association signals shared a consistent causal variant by reporting the posterior probabilities of following five hypotheses: H0, No association with either trait; H1, Association with trait 1, not with trait 2; H2, Association with trait 2, not with trait 1; H3, Association with trait 1 and trait 2, two independent SNPs; H4, Association with trait 1 and trait 2, one shared SNP. The results were denoted as PP0, PP1, PP2, PP3, PP4, higher PP4 suggested higher probability that a single variant effecting both two traits. After preparation of input files, we performed co-localization analysis using coloc.abf function with default setting. Genes of which the eQTL signal showed PP4 ≥ 0.7 with GWAS signal were called co-localized genes and considered as candidate OA risk genes.

To verify that our eQTL data have the capacity to nominate more effector genes for OA risk loci, we performed co-localization analysis using the eQTL signals of co-localized genes from all tissues in the GTEx project v8 cartilage eQTL data from ref.(10).

### Differential expression analyses

We performed differential expression analyses using the differential expression for repeated measures (dream) R package from variancePartition(86). For the comparison between damaged and intact cartilage, we produced the gene-level read counts matrix for 31 donors from whom we obtained paired damaged and intact cartilage samples. Genes with count per million (CPM) value < 0.1 in more than half samples were removed, then gene-level read counts were normalized with trimmed means of *m* values (TMM) method applied by edgeR (v3.40.2)(87) and voom-transformed with voomWithDreamWeights function from variancePartition R package. Besides cartilage type (damaged or intact), donor ID, sex, age, first two principal components of genotype, RNA integrity number (RIN), and sequencing batch were also included in the differential expression analysis model to correct the effects of technical and non-cartilage type-related biological factors. FDR ≤ 0.05 was considered as the threshold for the identification of differentially expressed genes.

### Processing of time-series gene expression data of chondrocytes differentiation

#### Soft-clustering of time-series expression data

To assess the functional roles of colocalized genes, we performed clustering and enrichment analysis on a chondrogenic differentiation time series RNA-seq data (GSE128554)(50). We supposed the genes with similar expression change patterns have similar functions. First, the gene-level read counts and TPM values of each sample were obtained with the same method used for quantification of genes of our cartilage samples. Then, pairwise differentially expressed gene identification was performed using R package DESeq2 (v1.28.1)(88). All genes with *P* ≤ 0.05 in at least one comparison were used for clustering analysis.

We performed soft-clustering of all differentially expressed genes based on the mean TPM values at each time point using R package Mfuzz (v2.48.0)(89). This method extracted the patterns of pre-specified *n* clusters and then assigned a membership score μ*_ij_* (0 < μ*_ij_* < 1) to gene *i* to indicate the degree of membership of this gene for cluster *j*. We chose to cluster the genes into 12 classes by visual observation. Finally, we considered genes with membership score ≥ 0.6 as final members for each cluster.

#### GO enrichment analyses

GO enrichment was performed by R package clusterProfiler (v3.16.4)(90) with default setting. GO terms with *P*-value ≤ 0.5 were considered statistically significant

### Cell culture

SW1353 cell line was obtained from Zeye (Shanghai, China). Cells were cultured in DMEM medium supplemented with 10% fetal bovine serum (Biological Industries, Israel), penicillin (100 U/ml) and streptomycin (0.1 mg/ml) in 5% CO_2_ at 37℃ incubator.

### Transfection and dual-luciferase reporter assays

A 876 bp putative open chromatin region containing rs11750646 (C alle or T alle), and a 1500 bp fragment surrounding *PI3KR1* transcription start site were inserted into pGL3 Basic vector (Promega, USA) (Supplementary Table 23). The luciferase reporter plasmid and phRLTK containing Renilla luciferase were co-transfected into cells using Lipo8000 Transfection Reagent (Beyotime, China).

After 48 h, the relative luciferase signal was computed as the ratio of firefly luciferase activity to Renilla signal, and relative activity was normalized by pGL3-promoter plasmid. All data came from four replicate wells, and statistical analyses were performed by a two-tailed *t* test.

### shRNA expression plasmid constructs and shRNA knockdown

For short hairpin RNA (shRNA) knockdown of *AR*, two independent oligonucleotides targeting *AR* were cloned into linearized miR30 backbone. The shRNA and negative control (NC) sequences are shown (Supplementary Table 23). A total of 2.5 μg of each plasmid were independently transfected into 80% confluence of SW1353 cells using Lipo8000 Transfection Reagent (Beyotime, China) according to the manufacturer’s instruction. After 48 hours of transfection, total RNA was isolated to detect the mRNA expression by quantitative real-time PCR (qRT-PCR). Moreover, *AR* shRNA plasmids were independently co-transfected with the expression plasmids including rs11750646 (C alle or T alle) with *PI3KR1* promoter used in the luciferase reporter assay. The transfection and measuring of luciferase activity are the same as indicated in dual-luciferase reporter assay section. All data came from at least three replicate wells, and statistical analyses were performed by a two-tailed *t* test.

### Cell line RNA isolation, reverse transcription, and quantitative real-time PCR

Total RNA was extracted using RNA fast200 Kit (Fastagen, China). 1 μg of total RNA was then reverse transcribed to cDNA using SuperScripts II First-Strand cDNA Synthesis Kit (Invitrogen, Waltham, MA). qRT-PCR was conducted with the QuantiTect SYBR® Green PCR Kit (Qiagen, Germany) on Bio-Rad instrument. The 2^−ΔΔCT^ method was used for data analysis. The qPCR primers are listed in Supplementary Table 23.

### Chromatin immunoprecipitation assay

Chromatin immunoprecipitation was performed using Simple-ChIP® Enzymatic Chromatin IP Kit (Magnetic Beads) (Cell Signaling Technology, USA). Pre-cleared chromatin was immunoprecipitated with monoclonal antibodies against AR (ab108341, Abcam, USA), normal rabbit IgG or antibody against acetyl Histone H3 (included in the kit). The total cell chromatin extract was used as input. Protein-DNA crosslinks were precipitated by using ChIP-Grade Protein G Magnetic Beads. And DNA is purified using DNA purification spin columns (included in the kit). The target DNA fragments were analyzed by Sanger sequencing of PCR fragments harboring SNP (Supplementary Table 23).

### Data visualization

ChIP-seq signals were visualized using R package Gviz (v1.32.0)(91). Heatmap was produced using R package ComplexHeatmap (v2.4.3)(92). Other results were visualized using R package ggplot2 (v3.3.6).

## Acknowledgements

We thank the patients who donated material for these studies. This work was supported by the National Natural Science Foundation of China (32170616, 32370653, and 82170896); Science Fund for Distinguished Young Scholars of Shaanxi Province (2021JC-02); Innovation Capability Support Program of Shaanxi Province (2022TD-44); Key Research and Development Project of Shaanxi Province (2022GXLH-01-22, 2023-YBSF-180), and the Fundamental Research Funds for the Central Universities. This study was also supported by the High-Performance Computing Platform and Instrument Analysis Center of Xi’an Jiaotong University.

## Author contributions

Conceptualization: T.-L.Y. and Y.G.; methodology: W.T., F.J., C.W., R.-H.H. and S.-S.D.; data analysis: W.T. and C.-Y.H.; experiment: C.W. and K.A.; visualization: W.T., C.W. and Q.-J.Z; resource: S.-Y.H., H.-M.S., H.-W.G.and Z.Y.; public data collection: W.T. and C.W.; writing-original draft: W.T. and C.W.; writing-review and editing: T.-L.Y. and Y.G.; project administration and supervision: T.-L.Y. and Y.G.

## Data and materials availability

Raw RNA-seq data reported in this paper have been deposited in the Genome Sequence Archive (GSA)(93) in National Genomics Data Center(94), China National Center for Bioinformation / Beijing Institute of Genomics, Chinese Academy of Sciences (GSA-Human: HRA006224) that are accessible at https://ngdc.cncb.ac.cn/gsa-human with reasonable request. Individual-level genotype data are available upon reasonable request: T.-L.Y. Access to individual-level genotypes from samples recruited within mainland China is subject to the policies and approvals from the Human Genetic Resource Administration, Ministry of Science and Technology of the People’s Republic of China. All lead variants of eQTL analyses (including regular eQTL analysis, conditional analysis, ci-eQTL analysis, and TFi-eQTL analyses), GWAS-eQTL co-localization, and allele-specific open chromatin results are presented as Supplementary Tables.

Processed knee cartilage single-cell RNA data can download from supplementary file at https://www.ncbi.nlm.nih.gov/geo/query/acc.cgi?acc=GSE104782. Knee cartilage eQTL summary data in ref.(10) is available at https://personal.broadinstitute.org/ryank/Southam_FunGen_eQTL_LowGradeCartilage.FastQTL_perm_nom_info.ForMSK-KP_16Jan2021.zip for low-grade (intact) cartilage and https://personal.broadinstitute.org/ryank/Southam_FunGen_eQTL_HighGradeCartilage.FastQTL_perm_nom_info.ForMSK-KP_16Jan2021.zip for high-grade (damaged) cartilage. The GTEx v8 eQTL summary data which contain associations of all tested variant-gene pairs is available at https://console.cloud.google.com/storage/browser/gtex-resources. Position Frequency Matrices (PFMs) of JASPAR 2022 core non-redundant motif set of vertebrates is available at https://jaspar.genereg.net/download/data/2022/CORE/JASPAR2022_CORE_vertebrates_non-redundant_pfms_jaspar.txt. Position Frequency Matrices (PFMs) of HOCOMOCO v11 motif set is available at https://hocomoco11.autosome.org/final_bundle/hocomoco11/core/HUMAN/mono/HOCOMOCOv11_core_HUMAN_mono_jaspar_format.txt. 1000 Genomes Project Phase 3 haplotype reference is available at https://mathgen.stats.ox.ac.uk/impute/1000GP_Phase3.tgz. Download links of GWAS summary data of osteoarthritis are presented in Supplementary Table 18.

Code used in this study is available at https://github.com/tianwen0003/OA-eQTL. The GTEx consortium modified version of FastQTL and corresponding auxiliary scripts were available at https://github.com/francois-a/fastqtl. Scripts used for gene-level filtration and PEER factor calculation are available at https://github.com/broadinstitute/gtex-pipeline/archive/refs/tags/gtex_v8.tar.gz. Pipeline code for mashR is available at https://github.com/stephenslab/gtexresults. Additional code may be available upon reasonable request from the corresponding authors.

