## Supplementary Figures 1-20 for "Comprehensive profiling of transcriptional regulation in cartilage reveals pathogenesis of osteoarthritis"

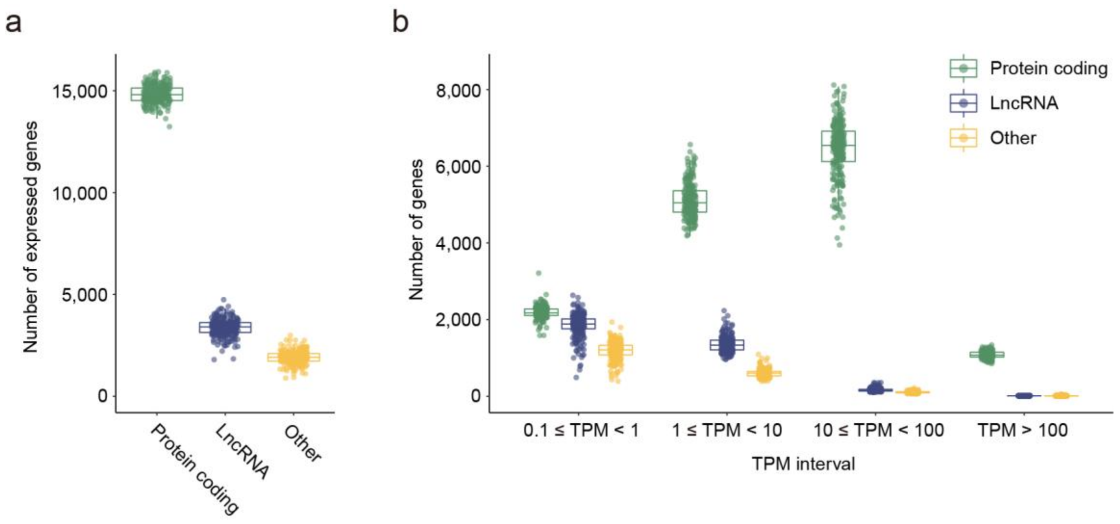


**Supplementary Figure 1. Summary of expression data. a**, Number of expressed (TPM ≥ 0.1) protein coding genes, long non-coding genes (lncRNA) and genes of other types per sample. Each dot is a different sample (*n* = 245). The x axis is different gene type; The y axis is number of expressed genes (TPM ≥ 0.1). **b**, Number of protein coding genes, long non-coding genes (lncRNA) and genes of other types in each expression value interval per sample. Each dot is a different sample (*n* = 245). The x axis is different expression interval; The y axis is the number of genes.

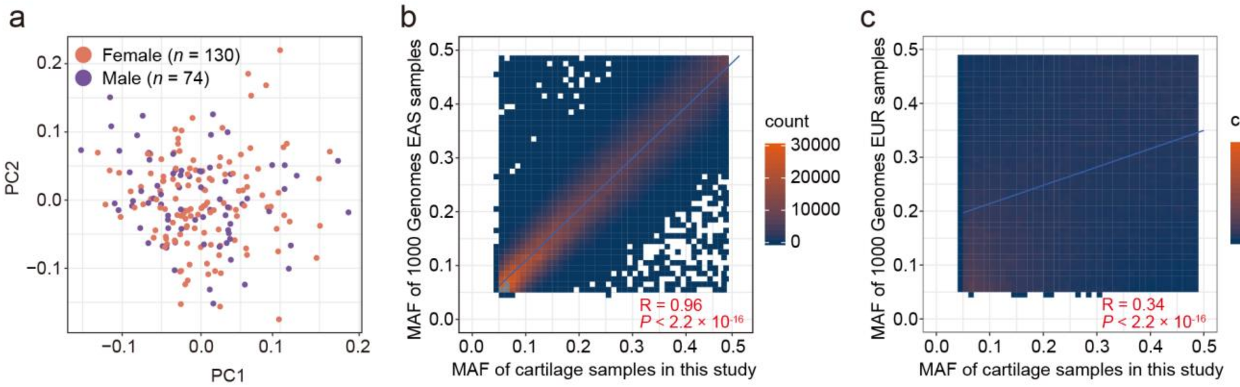


**Supplementary Figure 2 Summary of genotype data.** **a**, Principal component analysis (PCA) dimensionality reduction plot of the 204 genotypes data included in this study. **b**, Comparison of minor allele frequency (MAF) of all variants contained after imputation quality control between our samples and Asian populations in the 1000 Genomes Project. R is the Pearson correlation. The x axis it the MAF of variants in our study; The y axis is the MAF of variants in samples of Asian population in the 1000 Genomes Project. **c**, Comparison of the correlation of MAF of all variants contained after imputation quality control between our samples and European population in the 1000 Genomes Project. R is the Pearson correlation. The x axis it the MAF of variants in our study; The y axis is the MAF of variants in samples of European population in the 1000 Genomes Project.

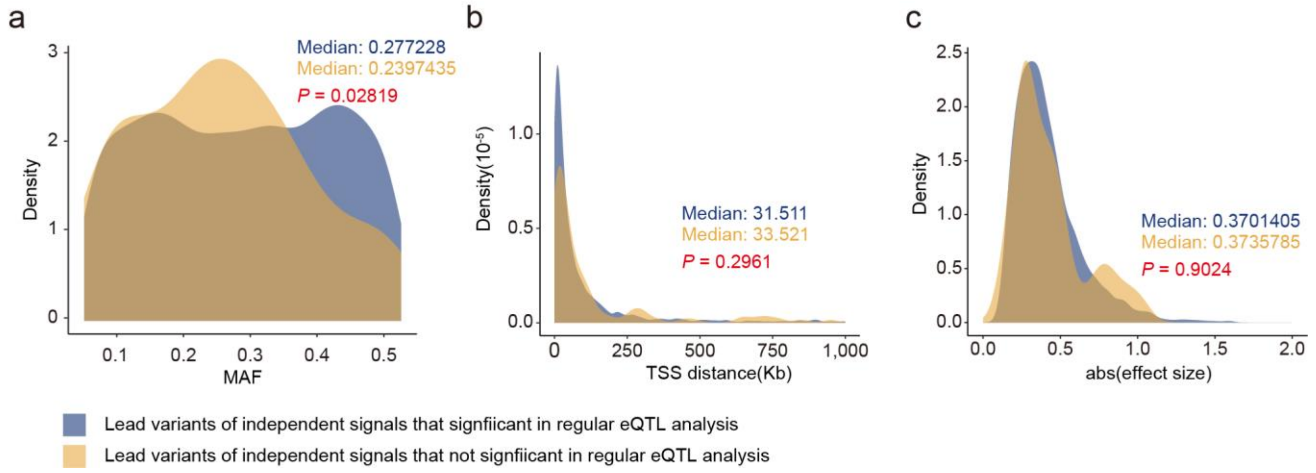


**Supplementary Figure 3.** Comparison of MAF (a), distance to transcription start site (TSS) (b), and effect size (c) of independent lead variants that originally significant in regular *cis*-eQTL analysis and that newly identified in conditional analysis. All *P*-values are calculated using Wilcoxon rank sum test.

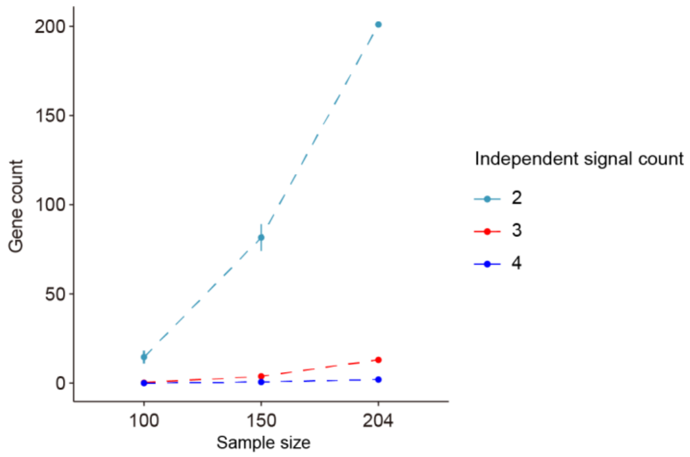


**Supplementary Figure 4. Downsampling for conditional analyses.** We take samples with size of 100 and 150 for reanalysis of regular *cis*-eQTL, and then get the number of genes with certain number of independent eQTL signals. Five dependent replications for each sample size (except 204). Data is showed as mean ± sd.

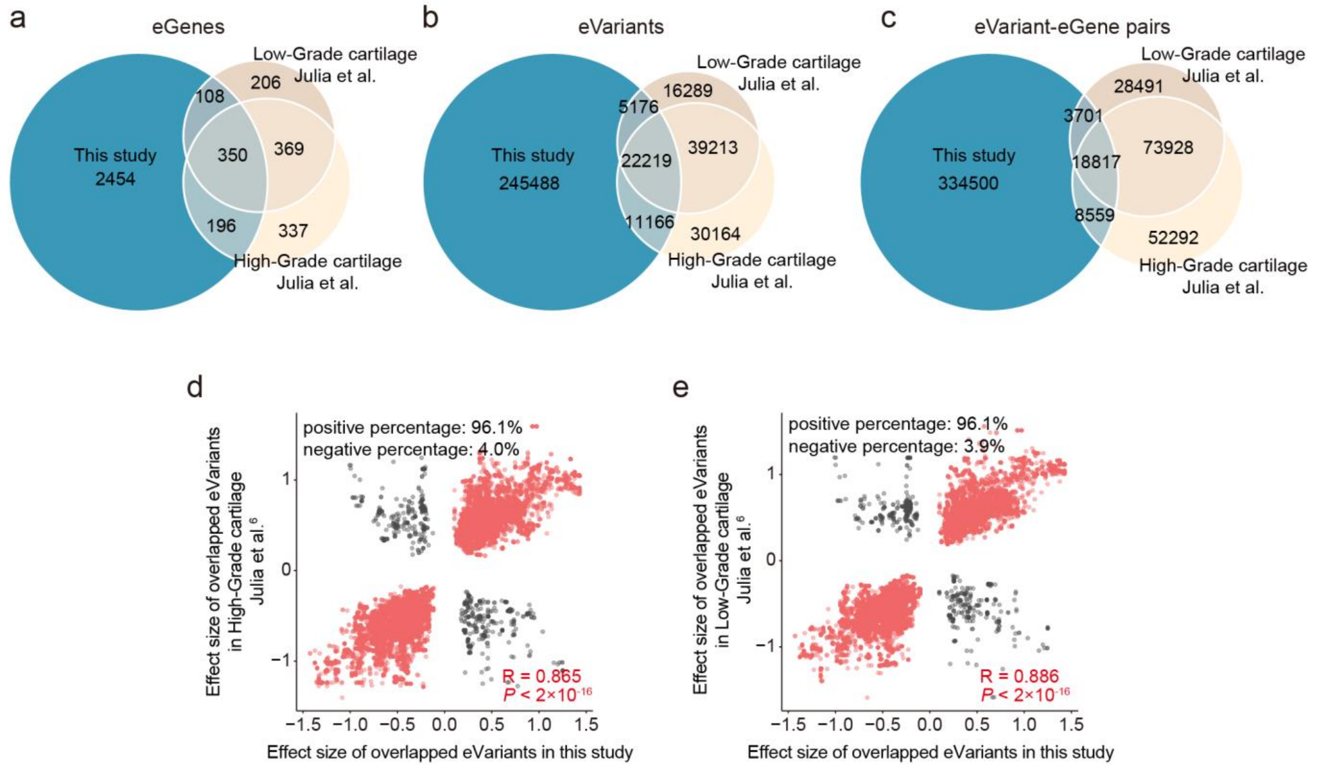


**Supplementary Figure 5. eQTL results comparison between datasets. a-c**, Overlap of significant eGenes (a), eVariants (b), eVariant-eGene pairs (c) between our study and published eQTL datasets of low-grade (intact) and high-grade (damaged) knee cartilage samples, respectively. **d,e**, Effect size comparison of the overlapped eVariant-eGene pairs between our study and published eQTL dataset of high-grade (d) and low-grade (e) cartilage samples. R is the Pearson correlation.

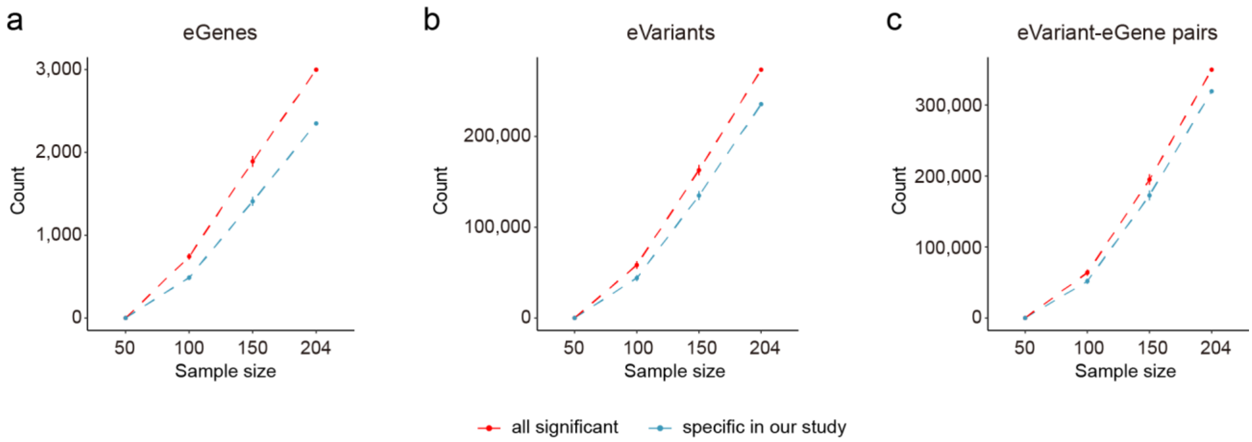


**Supplementary Figure 6. Downsampling for regular *cis*-eQTL analyses.** We take samples with size of 50, 100 and 150 for reanalysis of regular *cis*-eQTL, and then compared the results of eGenes (a), eVariants (b) and eVariant-eGene pairs (c) with merged published eQTL datasets of low-grade (intact) and high-grade (damaged) knee cartilage samples. Five dependent replications for each sample size. Data is showed as mean ± sd.

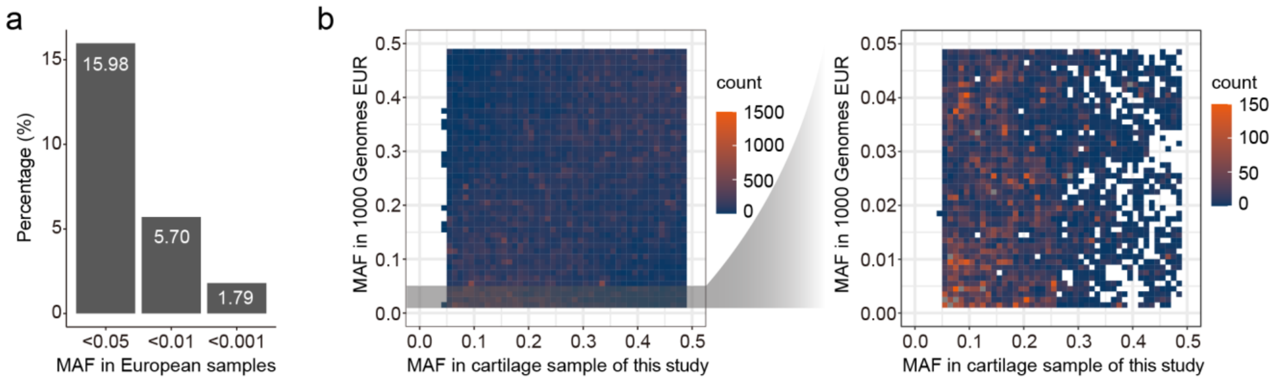


**Supplementary Figure 7. Comparison of minor allele frequency of our specific eVariants between Chinese samples and European samples. a**, Percentage of BIGC specific eVariants with minor allele frequency (MAF) less than 0.05, 0.01 and 0.001 in European samples of 1000 genome project. **b**, Comparison of MAF of BIGC specific eVariants between our samples and European samples in 1000 genome project.

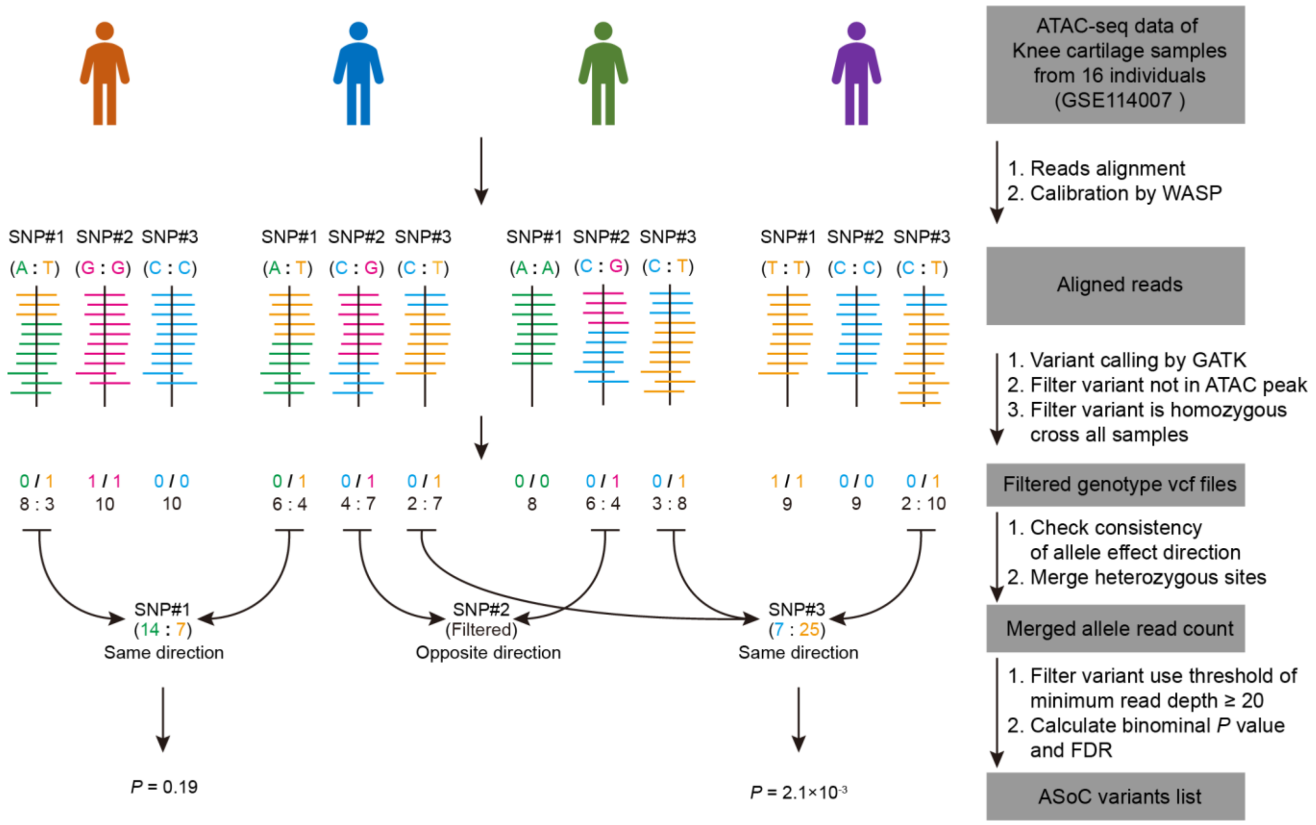


**Supplementary Figure 8.** Method for identification of ASoC variants. A total of 16 ATAC-seq data were used for analysis, here, we only showed four data for simplicity.

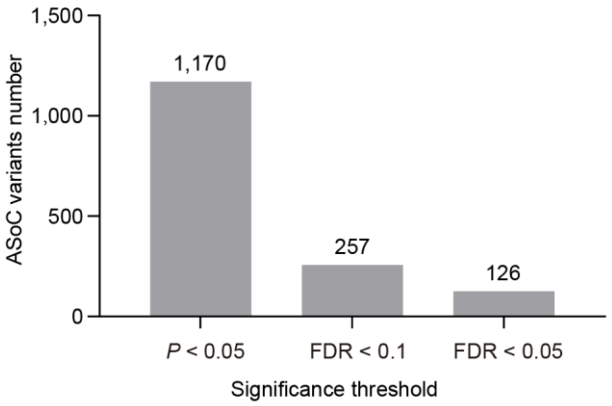

**Supplementary Figure 9.** Number of allele specific open chromatin (ASoC) variants identified using different significance threshold.

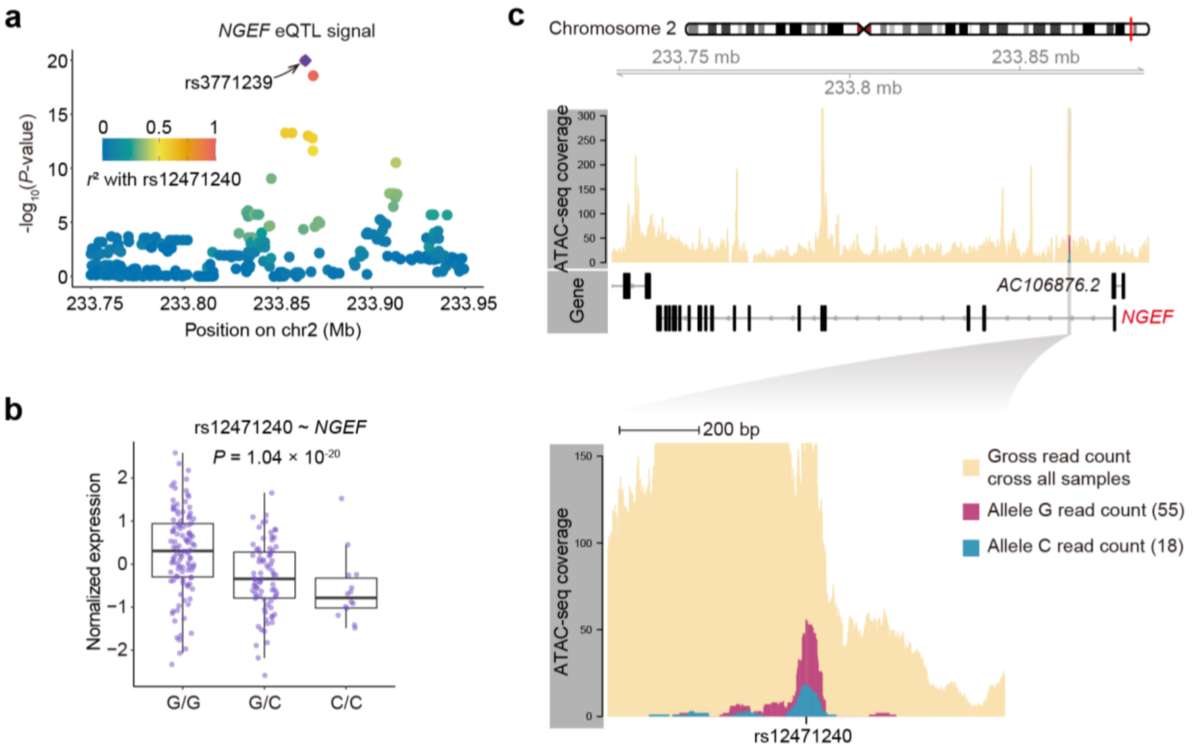


**Supplementary Figure 10. a**, Plot show eQTL signal associated to *NGEF*. The y axis is the statistical significance of cartilage eQTL for *NGEF* gene in -$\log_{10}$(*P*-value). The x axis is the chromosome coordinate. **b**, Association between the genotypes of variant rs12471240 and *NGEF* gene expression. The y axis is the normalized expression of *NGEF* in damaged cartilage samples; the x axis is the genotype at the rs12471240. The whiskers extend to the 5th and 95th percentiles; Each dot is a different sample. The *P*-value was reported by FastQTL nominal procedure. **c**, Variant rs12471240 located within an open chromatin region marked by an ATAC-seq peak in the intron region of the *NGEF* and altered the local chromatin accessibility. The sequencing reads with G allele piled up a significantly higher peak than that piled up by C allele-contained reads.

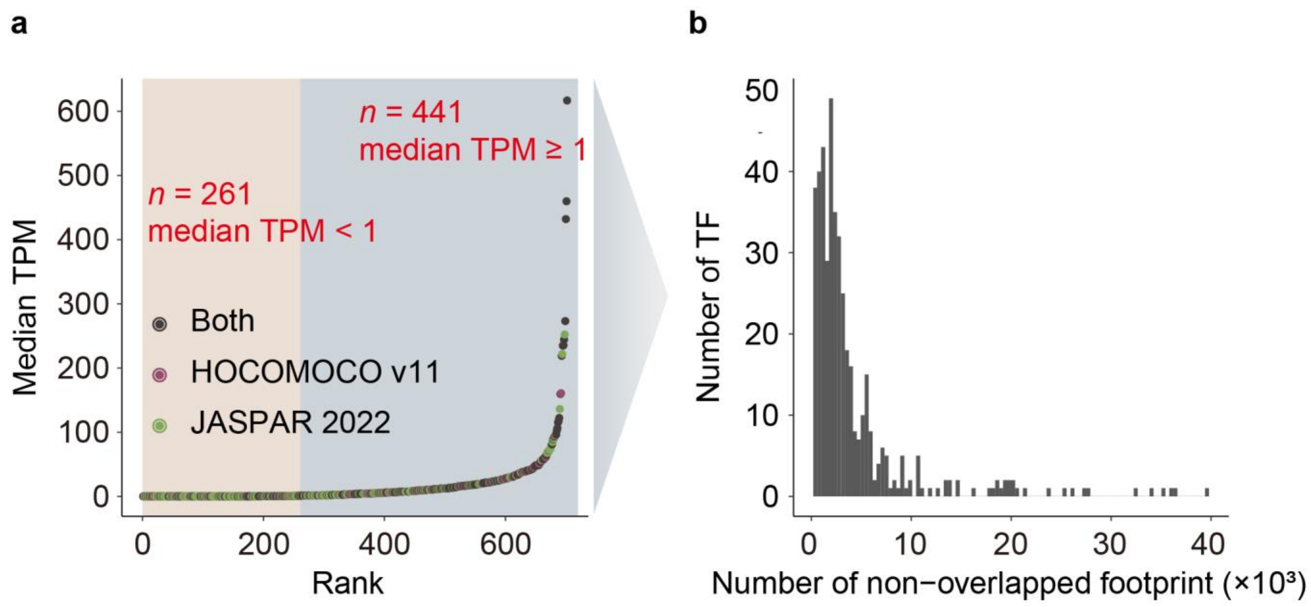


**Supplementary Figure 11. a**, Median expression across all damaged cartilage samples of TFs that have at least one motif in JASPAR 2022 core non-redundant or HOCOMOCO v11 motif sets. Each dot is a difference TF. The dots in dark grey represent TFs with motif in both two datasets; The dots in purple represent TFs only with motif in HOCOMOCO v11 motif set; The dost in green represent TFs only with motif in JASPAR 2022 core non-redundant motif set. The y axis is the median expression across all damaged cartilage samples of each TF; The x axis is the rank of median expression value of each TF in ascending order. **b**, Distribution of number of TFs with different number of footprints identified in all. The y axis is the number of TF; The x axis is the number of non-overlapped footprint.

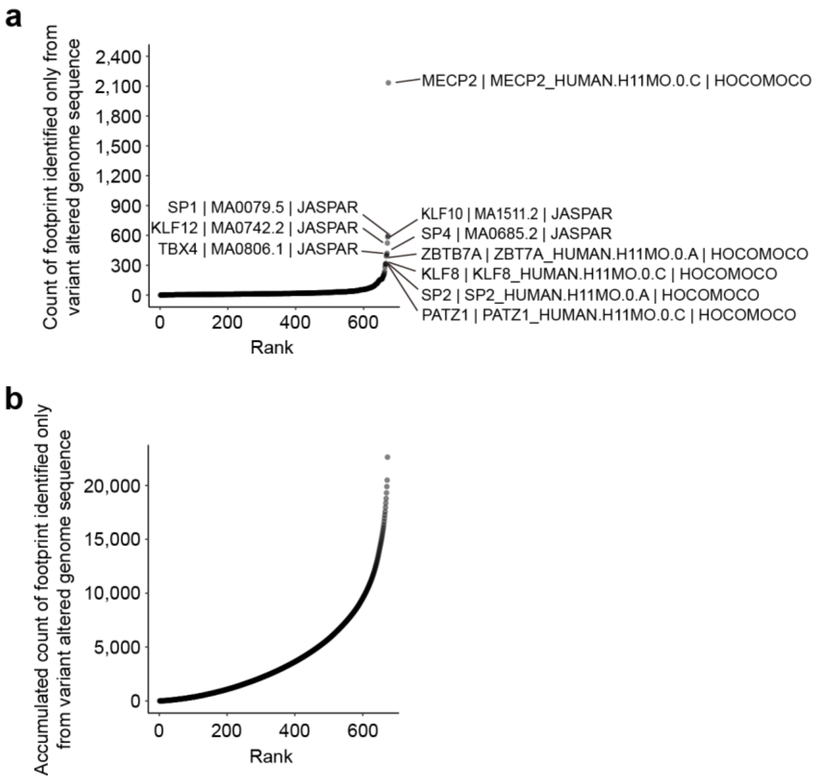


**Supplementary Figure 12. Allele sensitive footprint analysis contributes to the identification of more footprints. a**, Number of additional footprints identified for each motif with allele sensitive method. Each dot is a difference motif. The annotation text of each point is TF symbol, motif ID and dataset belong to of the motif, separated with short vertical line. The y axis is the number of additional footprints identified; The x axis is the rank of additionally identified footprints number for each motif in ascending order. **b**, Accumulated number of additional footprints identified for each motif with allele sensitive method. Each dot is a difference motif. The y axis is the accumulated number of additional footprints identified; The x axis is the rank of additionally identified footprints number for each motif in ascending order.

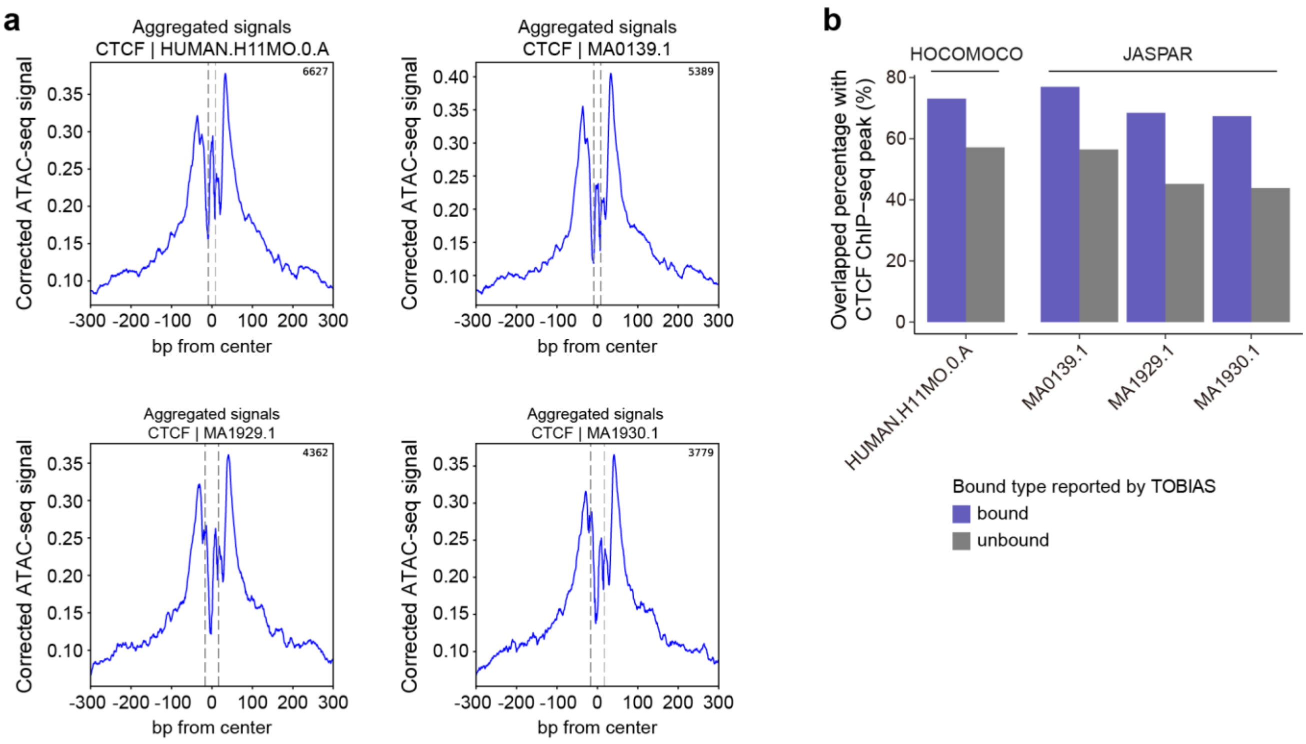


**Supplementary Figure 13. TF footprints validation by CTCF ChIP-seq data. a**, Aggregated ATAC-seq signal around four footprint sets corresponding to different CTCF motifs. The dashed lines indicate motif boundaries. **b**, Overlap percentage (%) of TOBIAS reported bound or unbound regions of CTCF with chondrocytes CTCF ChIP-seq peaks.

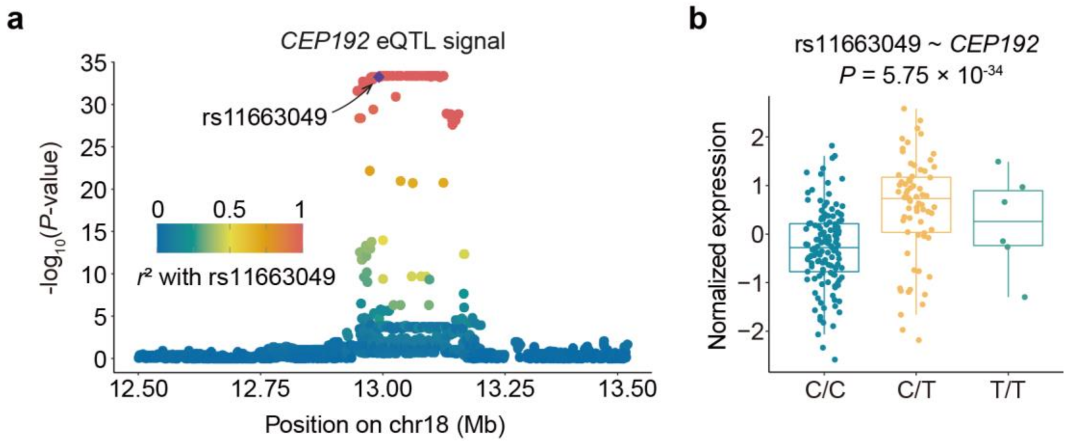


**Supplementary Figure 14. a**, Plot show eQTL signal associated to *CEP192*. The y axis is the statistical significance of cartilage eQTL for *CEP192*gene in -$\log_{10}$(*P*-value). The x axis is the chromosome coordinate. **b**, Association between the genotypes of variant rs11663049 and *NGEF* gene expression. The y axis is the normalized expression of *CEP192* in damaged cartilage samples; the x axis is the genotype at the rs11663049. The whiskers extend to the 5th and 95th percentiles; Each dot is a different sample. The *P*-value was reported by FastQTL nominal procedure.

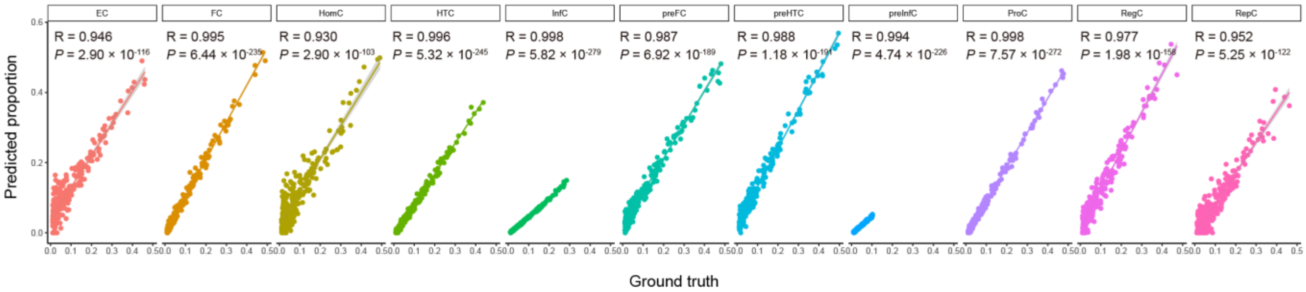


**Supplementary Figure 15. Signature matrix validation using pseudo-bulk express data produced with knee cartilage scRNA-seq data (GSE255460).** Each pseudo-bulk expression data was produced using 5,000 randomly selected cells from the cell list that not be used for signature matrix generation. Each dot is a different pseudo-bulk sample, 235pseudo-bulk samples produced in total. The x axis is the ground truth of cell proportion of pseudo-bulk samples, the y axis is the predicted cell proportions of pseudo-bulk samples using the signature matrix produce using CIBERSORTx with 5,000 randomly selected single cells

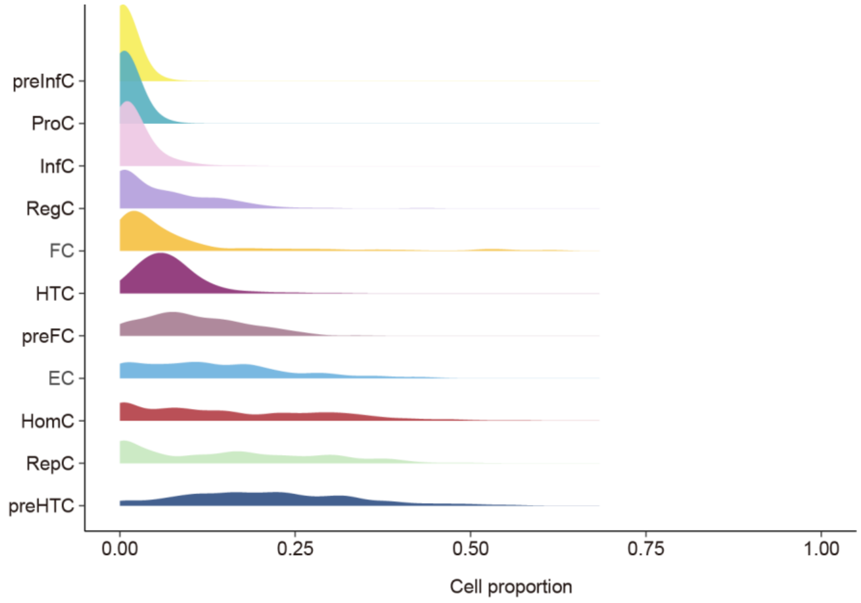


**Supplementary Figure 16.** Distribution of proportions of each chondrocyte subtype across all damaged cartilage samples. The y axis is the sample density; The x axis is the cell proportion.

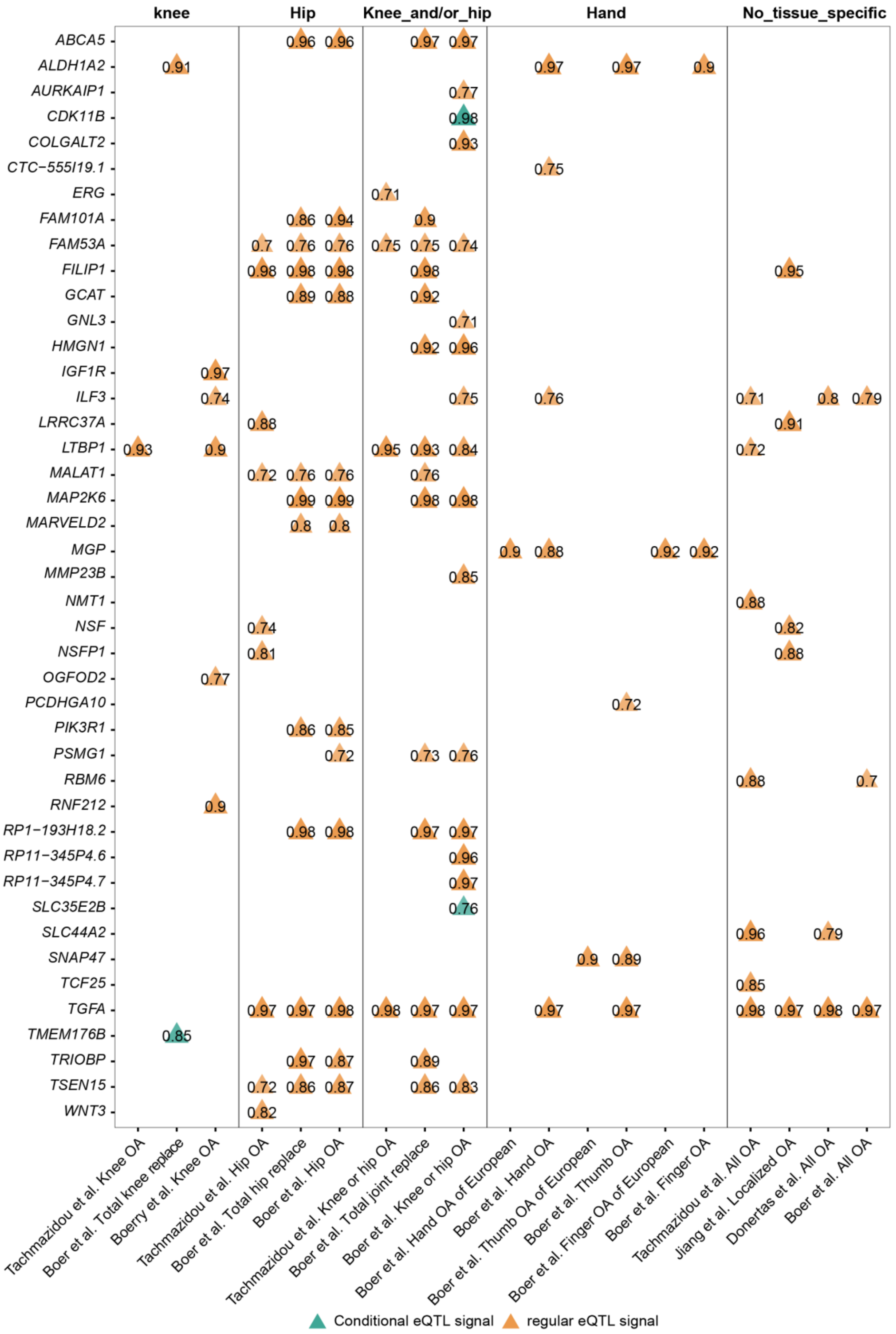


**Supplementary Figure 17.** Detailed co-localization results of OA GWAS loci with cartilage eQTLs. The numbers on the triangles represent the probability of a single shared variant for both GWAS and eQTL signal (PP4). Genes with PP4 ≥ 0.7 in at least one GWAS data were showed.

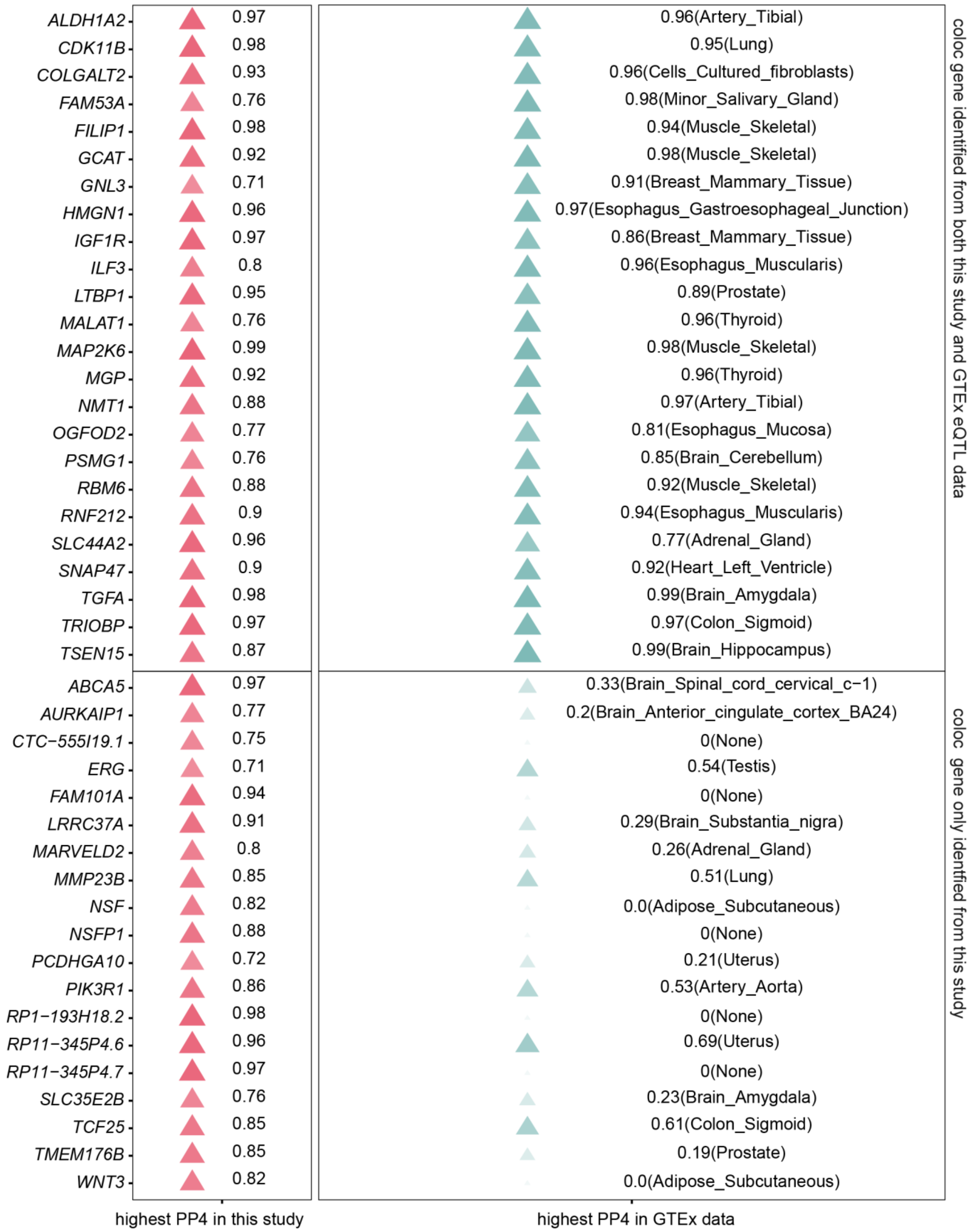


**Supplementary Figure 18.** Comparison of co-localization results when using our cartilage eQTL dataset and eQTL datasets of other tissues from the GTEx project. Tissue in parentheses represents the tissue from which the highest PP4 was obtained. Twenty-five genes were nominated with at least one eQTL datasets in the GTEx project at the threshold PP4 $\geq$ 0.7.

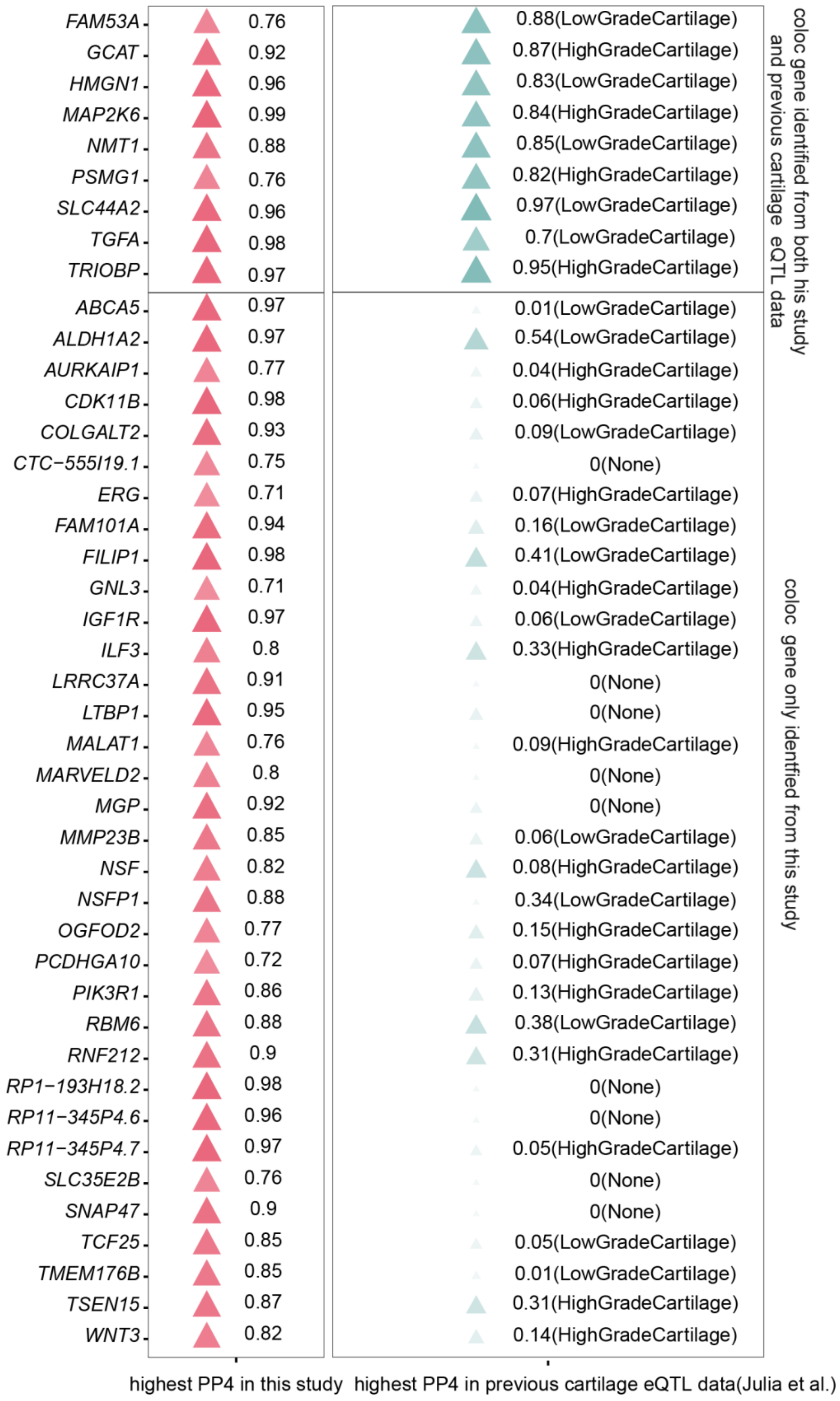


**Supplementary Figure 19.** Comparison of co-localization results when using cartilage eQTL dataset from our study and that from Julia et al. Tissue in parentheses represents the tissue from which the highest PP4 was obtained. Nine genes were nominated with cartilage eQTL datasets from Julia et al. at the threshold PP4 ≥ 0.7.

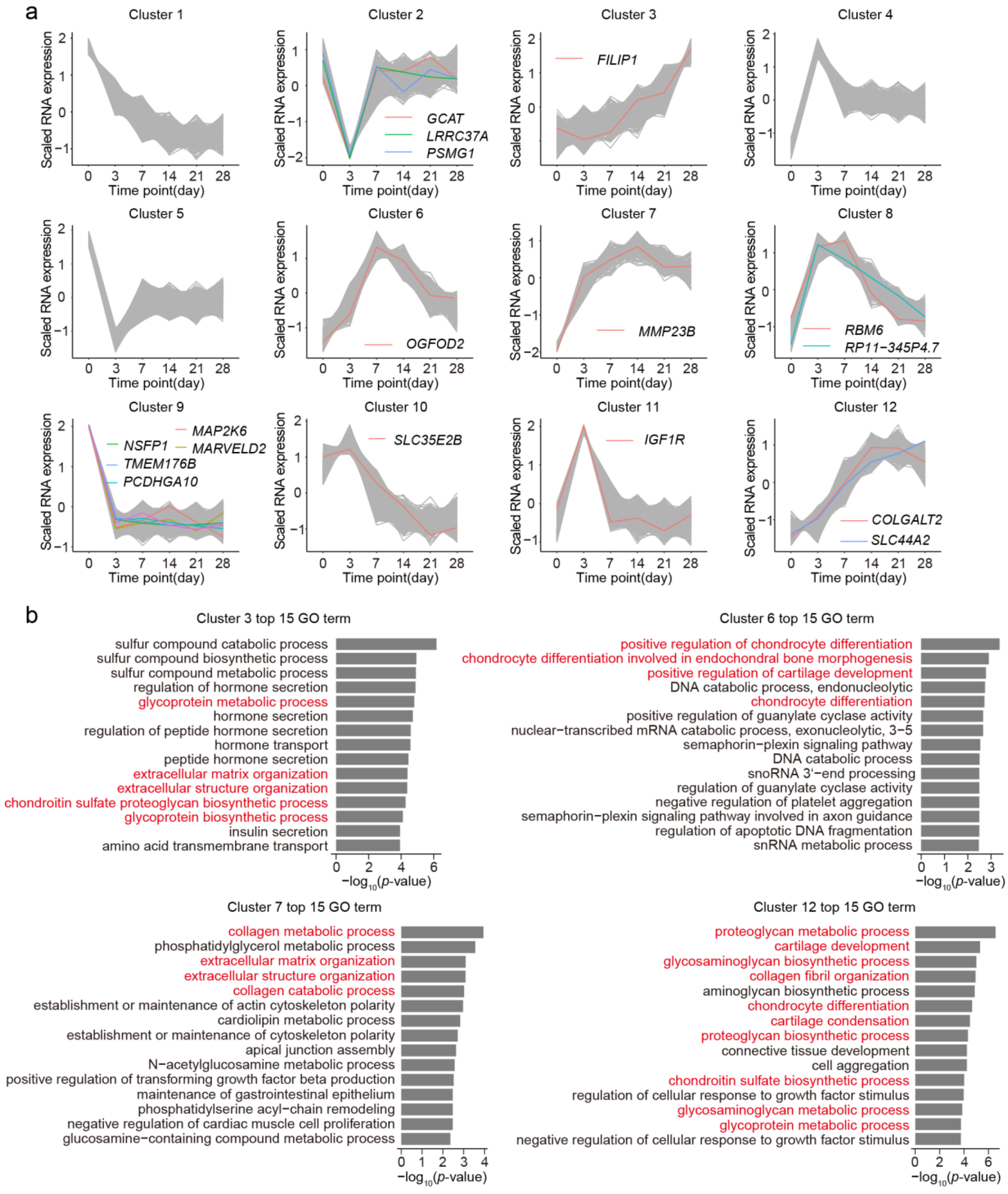


**Supplementary Figure 20. Generation of gene sets for OA risk genes annotation. a**, Soft-clustering results of time-series gene expression data of chondrocyte differentiation. The y axis is the scaled expression; The x axis is the time point. The co-localization analysis nominated risk genes belong to each cluster are shown by difference color. **b**, A total of four clusters of genes for which the top 15 enriched GO terms related to chondrocytes function. The y axis is enriched GO terms; the x axis is the enrichment statistical significance in -$\log_{10}$(*P*-value).
